# Reinforcement Learning for Bio-Retrosynthesis

**DOI:** 10.1101/800474

**Authors:** Mathilde Koch, Thomas Duigou, Jean-Loup Faulon

## Abstract

Metabolic engineering aims to produce chemicals of interest from living organisms, to advance towards greener chemistry. Despite efforts, the research and development process is still long and costly and efficient computational design tools are required to explore the chemical biosynthetic space. Here, we propose to explore the bio-retrosynthesis space using an Artificial Intelligence based approach relying on the Monte Carlo Tree Search reinforcement learning method, guided by chemical similarity. We implement this method in RetroPath RL, an open-source and modular command line tool. We validate it on a golden dataset of 20 manually curated experimental pathways as well as on a larger dataset of 152 successful metabolic engineering projects. Moreover, we provide a novel feature, that suggests potential media supplements to complement the enzymatic synthesis plan.

## Introduction

Efficient computational tools are required for metabolic engineering to achieve its true potential as a game-changer in the bioeconomy. Such tools include pathway design software, to assist the metabolic engineer in finding new pathways for production of valuable chemicals. While some tools restrict themselves to reactions already present in databases (Rodrigo et al. 2008; Moriya et al. 2010), others allow the generation of de novo reactions, using retrosynthesis algorithms (Henry, Broadbelt, and Hatzimanikatis 2010; Carbonell et al. 2011; Yim et al. 2011; Carbonell et al. 2014; Liu et al. 2014; Campodonico et al. 2014; Hadadi et al. 2016; Kumar et al. 2018; Delépine et al. 2018). At its core, a retrosynthesis algorithm is simple: break down a target molecule into simpler molecules that can be combined chemically or enzymatically to produce it and iterate recursively until all required compounds are either commercially available or present in the microbial strain of choice. Current bio-retrosynthesis tools suffer from several limits. First of all, they are usually accessible through a web-server and not open-source, limiting an expert user’s capacity to install them locally and tune them. Secondly, a number of parameters are often included within the pathway search, decided by the software designer and with limited capacity for a user to incorporate his own knowledge, solving for both retrosynthesis and parameters optimisation. Some examples include enzyme performance (Delépine et al. 2018), predicted yield (Campodonico et al. 2014; Carbonell et al. 2014; Liu et al. 2014; Cho et al. 2010; Tokic et al. 2018), thermodynamics or cofactor usage (Kumar et al. 2018). Moreover, those tools do not include the latest advances in combinatorial search space exploration, pioneered in the field of Artificial Intelligence.

To address those limitations, we propose to make use of the Monte Carlo Tree Search (MCTS) reinforcement learning algorithm which has already revolutionised the field of Artificial Intelligence, as illustrated by the stunning victory of Google’s AI (AlphaGo) against a Go master in 2016 (Silver et al. 2016, 2017, 2018). An interesting application used this algorithm combined with neural networks in chemical retrosynthesis, but acknowledging that natural compounds synthesis was beyond their scope (Segler, Preuss, and Waller 2018). To make available the algorithm to the wider audience, we implement it as RetroPath RL, an open source python package freely available on GitHub. RetroPath RL provides a programmatic access to discover biosynthesis pathways using MCTS, while allowing a number of augmentations and features to be used or developed by expert users to tune it to their needs. Lack of open-source computer-assisted pathway design tools is currently pointed out as one of the major limits faced by metabolic engineering projects (Casini et al. 2018; Lee et al. 2019). Therefore, RetroPath RL is a timely software that will hopefully contribute to metabolic engineering realising its true potential for green chemistry.

We propose below an explanation of the theoretical background giving context to the present work.

### Reaction rules for representing enzymatic reactions

Reaction rules describe the changes in bonding patterns when a set of substrates is transformed into a set of products in order to encode enzymatic reactions for retrosynthesis. An important feature of rules for retrosynthetic applications is that they need to be generalisable, so that they can be applied to a new substrate that was not from amongst the substrates the rules were learned on. Moreover, using generalised reaction rules is the first step towards predicting promiscuous reactions, as those reactions are often missing from metabolic databases. Modelling promiscuity is a key feature in metabolic engineering, as it has for example been estimated that 37% of *Escherichia coli* K12 enzymes have a promiscuous activity on structurally similar substrates (Nam et al. 2012).

In our data-driven approach, we learn rules at various levels of specificity around the reaction present in the database, by keeping in the described pattern of the rule a varying number of atoms around the reaction centre. We select those atoms using a number we call diameter that represents the distance in bonds around the reaction centre: a rule at diameter 2 will include atoms at a distance of 1 around the reaction centre, while a rule a diameter 10 will include atoms at a distance of 5 around the reaction centre. Therefore, the rule at diameter 2 can apply to more diverse substrates and therefore encode more promiscuity than the rule at diameter 10. A more detailed description of reaction rules can be found in (Delépine et al. 2018; Duigou et al. 2019) and in the Methods section.

### Necessity of ranking reactions

The number of rules that can be applied greatly varies depending on the level of promiscuity considered and the chemical structure of the substrate. If this number is low enough, an exhaustive search can be considered, but otherwise, applying chemical reaction rules iteratively on substrates and their products leads to a combinatorial explosion. Encoding of reaction rules at various diameters allows selection of the degree of promiscuity, which has a high impact on those statistics. As an example, results for the rule sets used in this study are presented in Table 1, and for individual diameters in **Supplementary Table 1**. We can see from this Table that the branching factor (the average number of rules that apply to a substrate) is around 100 when using rules at diameters 6, 10 and 16 (slightly promiscuous, medium and very specific) and drastically increases with the level of promiscuity (adding rules at diameter 2 that are highly promiscuous lead to a branching factor of 900). Therefore, the more promiscuity we allow, the higher our branching factor becomes.

**Table 1:** Average number of applicable rules on a compound according to the set of diameters used. Information on average number of rules at all individual diameters are available in **Supplementary Table 1**.

| Set of diameters used | 6, 10, 16 | 2, 6, 10, 16 | 2, 4, 6, 8, 10, 12, 14, 16 |
| --- | --- | --- | --- |
| Average number of applicable rules | 100 | 898 | 1183 |

Such a high branching factor is comparable to the Go game (branching factor of 250) and much higher than chess (35), and the reason why using algorithms that were successful in this domain could also be of interest for retrosynthesis (Silver et al. 2016; Segler, Preuss, and Waller 2018). We therefore use an algorithm (MCTS) that can effectively handle this combinatorial explosion, and a heuristic (chemical similarity) to guide the search.

### Chemical similarity and sequence availability for reaction ranking

Chemical similarity between query (applied on a new substrate) and the native chemical transformation has been used in various studies (Cho et al. 2010; Hadadi et al. 2019; Coley et al. 2017; Campodonico et al. 2014). We adapted the strategy from Coley et al. that proceeds in a 2-step evaluation of the reaction. In a first step, before rule application, similarity between query and native substrates is calculated. After rule application, similarity between native and query products is also calculated. This allows accounting of similarity in a manner straightforward to use with mono-component reaction rules (Figure 1A). Using this metric allows us to select chemical reactions similar to the ones present in metabolic databases, increasing our chances that this predicted reaction can be catalysed.

**Figure 1:**
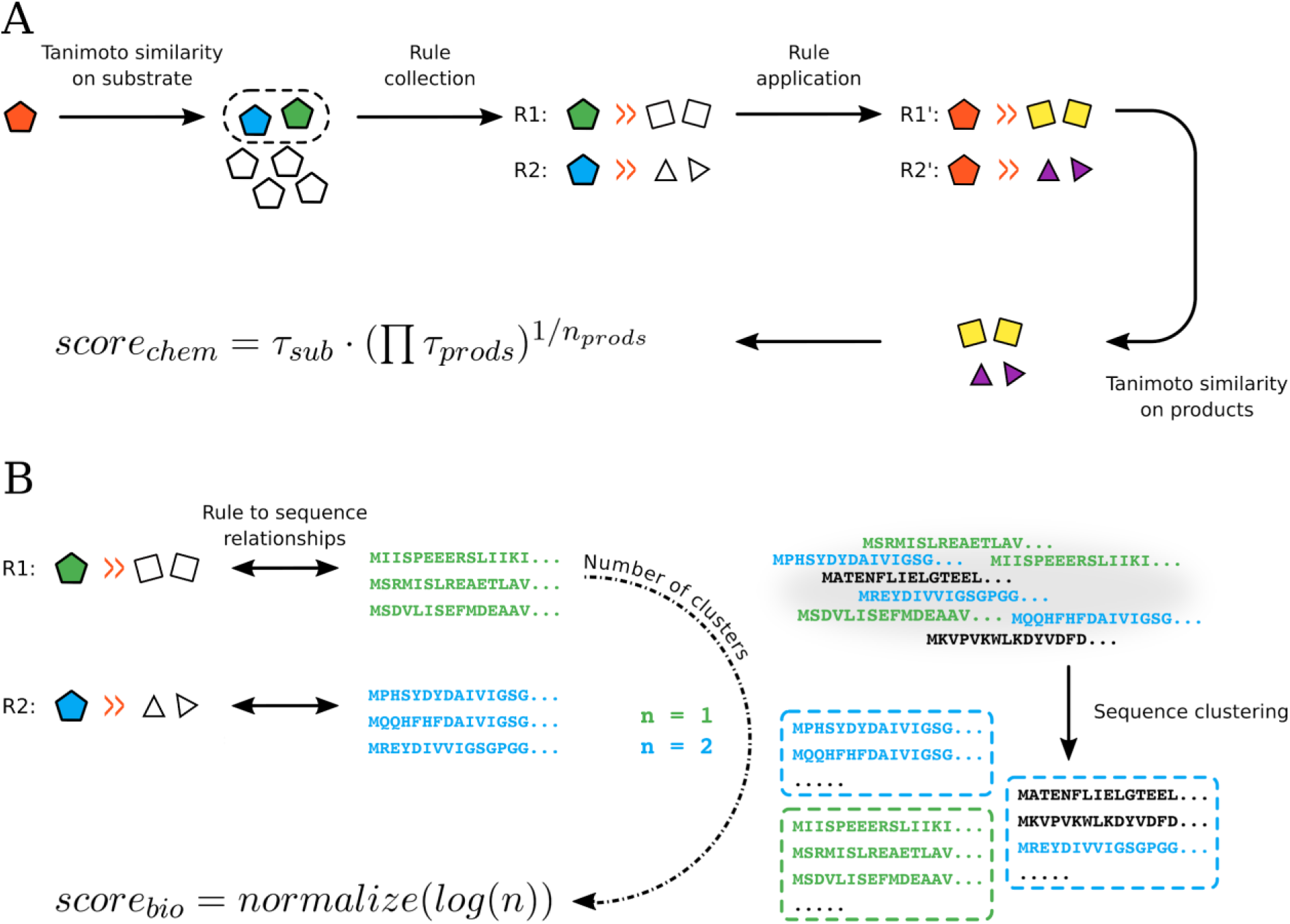
Chemical and biological score computations. For chemical score (**A**), we start by selecting substrates within the rule collection that are similar to the query substrate. We then apply those rule templates and check similarity of those products to the products the rules were learned on. For biological score (**B**), we establish rule-to-sequence relationships, and cluster all sequences based on sequence similarities. For each rule, we then count the number of cluster *n* spanning all related sequences. Score normalisation is detailed in Methods.

However, in (Cho et al. 2010; Hadadi et al. 2019; Coley et al. 2017), the authors are interested in chemical retrosynthesis, whereas we are interested in enzyme-catalysed reactions. Therefore, we combine (through multiplication) this chemical score with a scoring scheme that we developed previously (Delépine et al. 2018). Briefly, this biological score characterises our confidence that a sequence exists to catalyse the desired enzymatic rule. We updated this scoring scheme to be a normalised score, between 0 and 1, to be in the same range as the chemical score (Figure 1B). This biological score has the useful property that rules at low diameter (i.e. more promiscuous) are usually ranked lower than rules at high diameters (i.e. more specific and trustworthy).

In RetroPath RL, this combined biochemical score is used for ranking and excluding reaction rules that are not considered trustworthy (similarity too low to the original reaction, or sequence availability too low).

### Integrating rule ranking with the Monte Carlo Tree Search reinforcement learning method

Reinforcement learning methods are a class of machine learning algorithms that learn how to take actions in an environment so as to maximise a cumulative reward. Their main strength is their capacity to balance exploration of unchartered space and exploitation of current knowledge.

A Monte Carlo Tree Search is a reinforcement learning algorithm that explores the unknown space with random sampling. It proceeds in 4 phases, repeated until a resource budget (time or number of iterations) has been exhausted (Figure 2):

**Figure 2:**
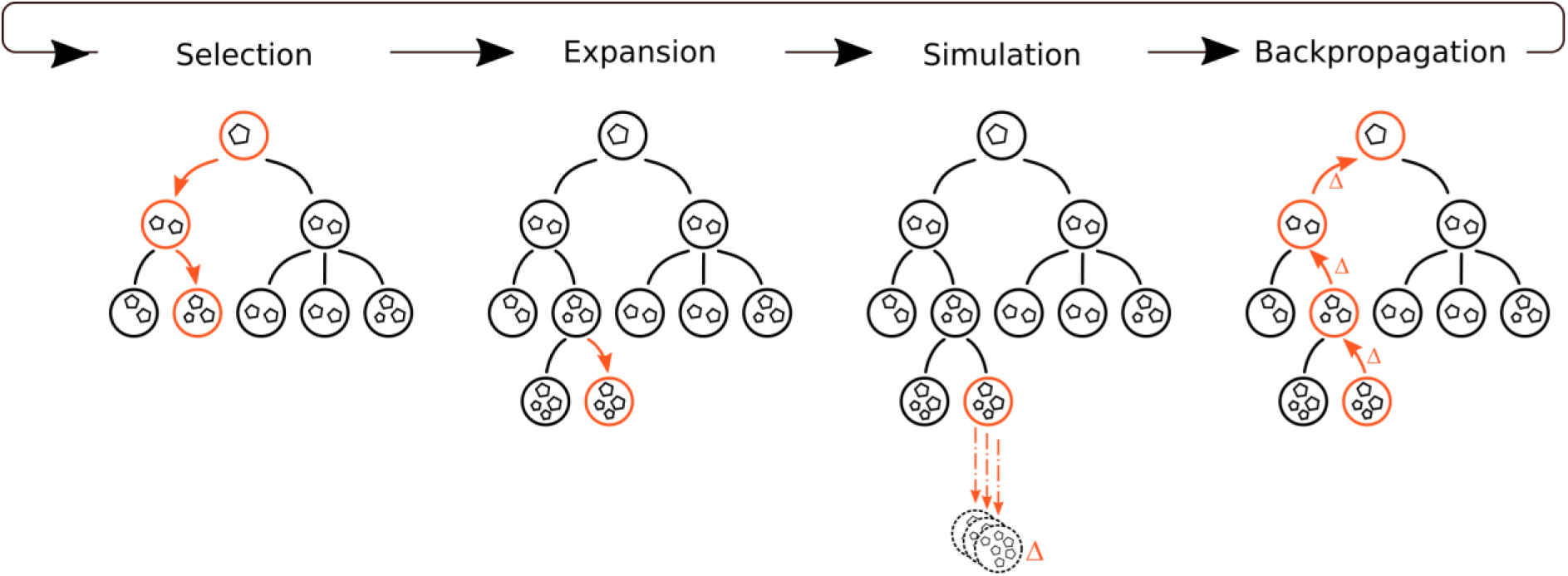
Monte Carlo Tree Search algorithm. Circles represent nodes, and pentagons molecules. Detailed explanations are in the main text and in the Methods section.

- **Selection.** Starting from the root node (here, a chemical state containing the target compound), the best child nodes according to the selection policy are iteratively chosen until a leaf node is reached. In the proposed implementation we use a selection policy guided by the biochemical score of the reaction rule, unless otherwise stated.
- **Expansion.** Possible transformations are selected based on the ranking scheme presented above and the node is expanded (with a predefined maximum number of children).
- **Simulation or rollout.** This is an iterative procedure that starts by checking the status of the state. If it is terminal, a reward (or penalty) is returned according to the rewarding policy (detailed in the methods section). If it is not terminal, a transformation is randomly sampled from available transformations and the process is repeated. This is performed until a maximum number of rollout steps or the maximal depth of the tree is reached.
- **Backpropagation or update.** The score obtained after exploring this node is returned to its parents to update their values and visit counts.

## Results and Discussion

### Evaluating RetroPath RL with a golden dataset

We first evaluate our implementation of MCTS for bio-retrosynthesis on a manually curated dataset of 20 compounds to identify the best settings for a retrosynthetic search on those compounds (chosen compounds and rationale for selection are available in the Methods section). This dataset allows us to verify bio-retrosynthesis tools suggest pathways that have been experimentally described in the literature, ensuring the biological relevance of the predicted pathways. Comparing RetroPath RL‘s features for expert users (see **Supplementary Note 1)** on this golden dataset allows us to select the best parameters possible for biological relevance. While various metrics could be available to describe what a good bio-retrosynthesis algorithm should do, there is no obvious consensus. Should such an algorithm be fast? Return a lot of pathways? Return fewer but more reliable pathways? We use three criteria for comparing algorithms. First, it should return pathways for as many compounds as possible. Second, results should include the chosen literature-described experimental pathway (exact intermediates are found). For parameter sets with identical results on those two criteria, the third criterion is that a better parameter combination should return the experimental pathway in fewer iterations.

The best parameters set we found used chemical and biological thresholds of 0.3, and a maximum of 10 allowed children per node (detailed configuration is available as **Supplementary Table 2** and effects of various parameters were investigated and presented in **Supplementary Note 1**). To generate the results presented in Figure 3, we ran RetroPath 2.0, and RetroPath RL with its default configuration and a configuration with more tolerant chemical scores for rule application. Detailed analysis of results on this dataset is available in **Supplementary Note 2**. We provide an example result in Figure 4 for mesaconic acid, interesting as it could be a renewable precursor to the commodity chemical (Yadav et al. 2018).

**Figure 3:**
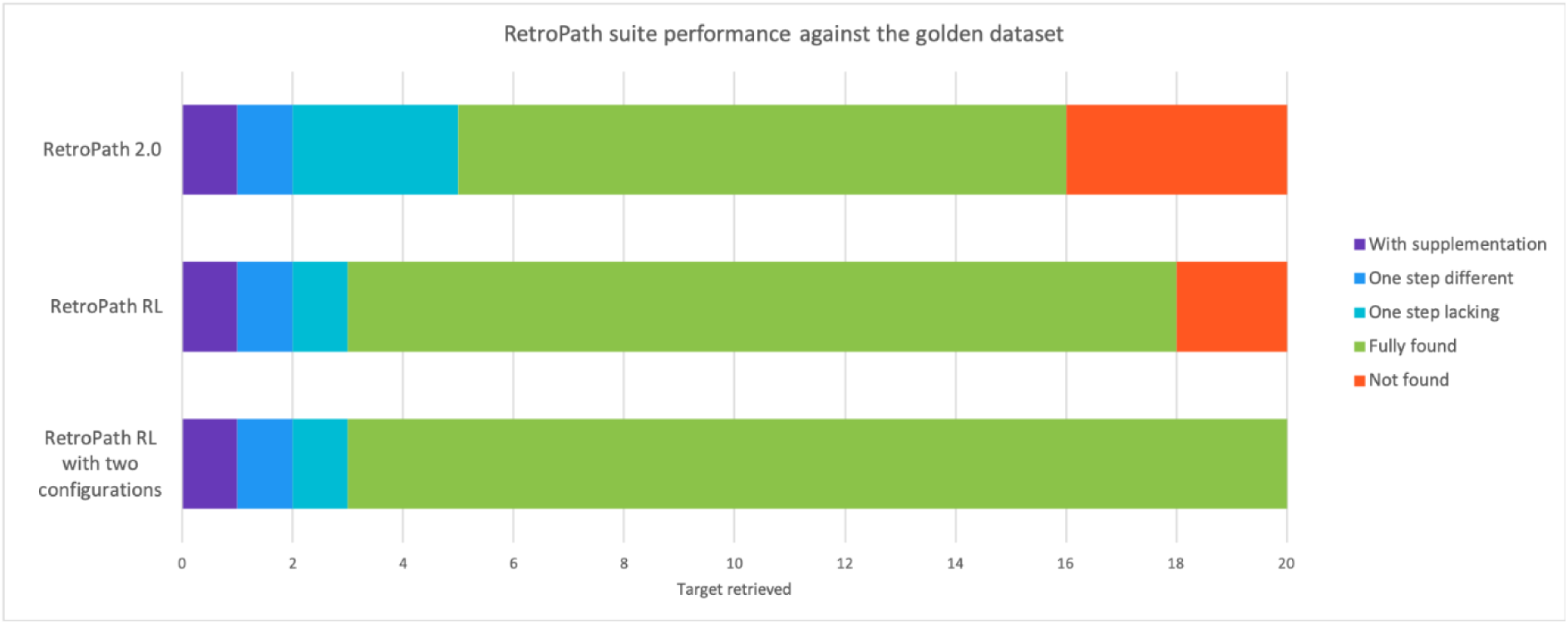
Results of the RetroPath suite against the golden dataset to identify the experimental pathway. We compared results of RetroPath 2.0, the default configuration of RetroPath RL and a combination of results between the default configuration and a more tolerant one on the used scores with a timeout of 3 hours. With supplementation (purple) means a supplement has to be provided in the media to identify the correct experimental pathway. One step different (dark blue) means only one step differs from the described pathway, for example by using a different co-substrate. One step lacking (light blue) means the search algorithm found a pathway identical to the experimental one, except one step which was short-cut. Fully found (green) means the experimental pathway was found without restriction. Not found (orange) means the experimental pathway was not found.

**Figure 4:**
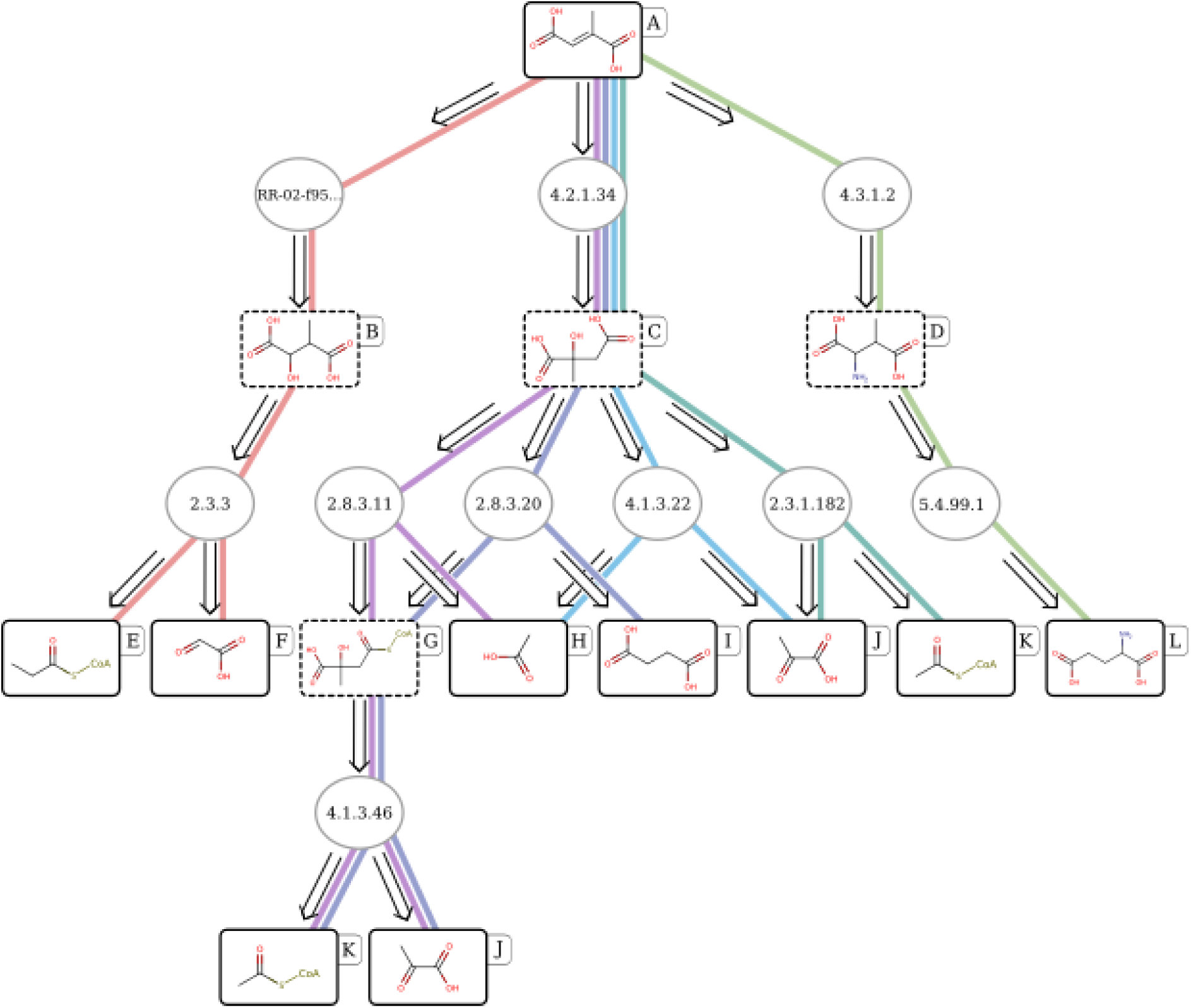
Example of bio-retrosynthetic scope obtained from RetroPath RL for mesaconic acid. Dead end compounds have been removed, pathways lying in the scope are depicted with distinct colours. Compounds are represented by their chemical structures and reactions by their EC numbers or their rule ID of no EC is known. Mesaconic acid (at the top) and sink compounds are surrounded by a solid line, while intermediates are surrounded by a dashed line. Only the pathways predicted up to 3 steps are shown. Compound names: mesaconic acid (A), 3-methylmalic acid (B), citramalic acid (C), 3-methylaspartic acid (D), propanoyl-CoA (E), glyoxylate (F), citramalyl-CoA (G), acetate (H), succinate (I), pyruvate (J), acetyl-CoA (K), glutamate (L).

We can see from these results that the algorithm encoded within RetroPath RL suggests experimentally relevant pathways for biochemists, as it finds the exact pathway described in the literature 75% of the time with strict settings, and 95% of the time when trying more tolerant settings on failed compounds, media supplementation or using another cofactor (lacking one step is considered a failure). We can see this is better than our previous algorithm, proving that RetroPath RL suggests more experimentally relevant pathways for metabolic engineering.

### Importance of our scoring schemes

Although all parameters of interest are evaluated in **Supplementary Figures 1 to 12** in **Supplementary Note 1**, we detail here the impact of the scoring cut-offs and scheme. As mentioned in the theoretical background section, we use a biochemical score, based on both chemical similarity and estimation of enzyme sequence availability. We analysed algorithm behaviour for various biological, chemical and biochemical scores, as well as when guided only by similarity, biological score or no scoring scheme (classical algorithm). The results are presented in Figure 5 and validate our approach. We can see the score contributing most to the importance of the cut-off parameter values is the biological score that ensures experimental availability of the enzymatic sequence, while the score that contributes most to the guidance of the Monte Carlo Tree Search is the chemical similarity component, showing both scores are necessary for ensuring efficient search and experimental relevance of predictions.

**Figure 5:**
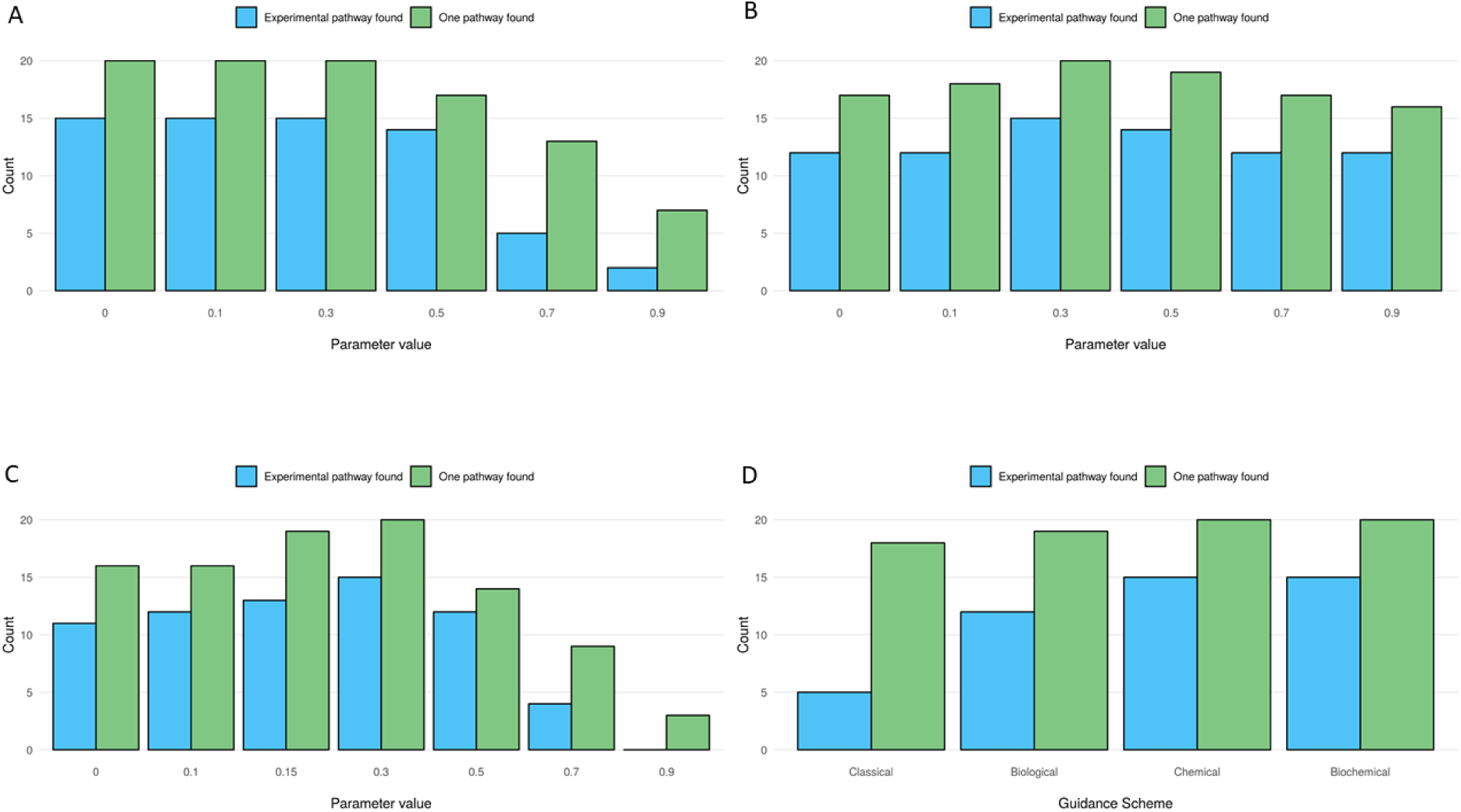
Impact of scoring scheme on retrieval performance of RetroPath RL. We compared results between using a biological score cut-off (**A**), a chemical score cut-off (**B**) and a biochemical score cut-off (**C**) varying between 0 and 0.9. In (**D)** we compared results between guiding the search based on the Classical UCT formula, a formula guided by Biological scoring, Chemical scoring or Biochemical scoring. One pathway found means that at least one pathway has been predicted. Experimental pathway found means that the experimental pathway is from amongst the predicted pathways.

### Evaluating RetroPath RL on successful metabolic engineering projects

After validating biological relevance of predictions for metabolic engineering, we tested RetroPath RL on a larger dataset. Our previous tool (Delépine et al. 2018) was tested on the LASER database (Winkler, Halweg-Edwards, and Gill 2015, 2016) that compiles successful metabolic engineering projects, completed with compounds taken from the Metabolic Engineering journal (see Methods, available as **Supplementary Data 1**). Given the curation level of this dataset, we checked the number of compounds for which we could find a pathway, and not the exact experimental pathway.

We ran our RetroPath 2.0 software using the same RetroRules rules (diameters 6, 10 and 16) and solved 77.6% compounds (118 out of 152), finding a median number of pathways of 4.5. With RetroPath RL and a score cut-off of 0.3, we solved 121 compounds (79.6%) with a median number of pathways of 11.5. Without score cut-off, we obtain 6 more compounds, yielding a success rate of 83.6% (results are available as **Supplementary Data 2**). The main advantage of the Monte Carlo Tree Search algorithm implemented in RetroPath RL over the brute force algorithm of RetroPath 2.0 is its capacity to find longer pathways. Indeed, the memory requirements of the exhaustive search performed by RetroPath 2.0 essentially limit it to 5 step pathways, while RetroPath RL can explore longer pathways and therefore find more solutions given the same allowed time. Its focus on promising areas of the search space also lead RetroPath RL to propose more solutions for the same compound.

### Supplement finder for media supplementation

The literature pathway that we identified for TPA used xylene for media supplementation, i.e. xylene is not a metabolite from the microbial strain but was necessary for producing TPA and experimentally added to the strain’s growth media. Knowing this allowed us to add xylene to the list of starting compounds for retrosynthetic search. Media supplementation is commonly done in metabolic engineering but rarely integrated into retrosynthesis tools. We therefore developed a RetroPath RL feature that analyses search trees and suggests potential media supplements, available from the chemical provider Sigma. We first extracted supplement information from LASER, as well as two TPA experimental pathways known from the literature. Filtering out gene inducers, antibiotics or early precursors, we obtain a list of 8 curated pathways with supplementation (available as **Supplementary Data 3**). In 5 cases out of 8, we retrieve the compound that was used for supplementation in the described experimental pathway. For 2 cases, we did not find the experimentally described supplement but suggest other possibilities. Here, we used availability in the Sigma-Aldrich catalogue as a criterion on whether a compound could be an interesting supplement. However, this feature can be used with any database of interest, for example with in-house compounds of a laboratory. One could include criteria such as capacity to cross membranes, solubility, toxicity, cost or any other feature of interest and select compounds that are biologically relevant for their application of interest.

### Custom use of RetroPath RL: avoid toxic intermediates

We sought to make our tool as modular and flexible as possible, therefore allowing expert users to input their knowledge. We showcase this by implementing a toxicity score to bias the search away from toxic compounds (Planson et al. 2012; Carbonell et al. 2014). This toxicity score is negative (between 0 and −10 in our training dataset) for toxic compounds and set to 0 otherwise. The strength of using bias in MCTS is that it favours preferred routes (here, avoiding toxic intermediates) but can still find results if those preferred routes are not successful, in contrary to other algorithms that would exclude those intermediates altogether. While implementing this feature did not change our results on the golden dataset (i.e. the experimental pathway was identified for the same compounds), the order as well as the total number of returned pathways was impacted (**Supplementary Table 3**).

While we tested it with toxicity, biasing the search could also be used to encourage pathways to be found from a set of privileged metabolites (core metabolism), by cost, availability in the cell or any other metric the advanced user wishes to use, making MCTS ideal to incorporate biological knowledge into retrosynthetic search.

## Conclusion and remarks

We showcase here the use of Monte Carlo Tree Search for bio-retrosynthesis, and provide an implementation as RetroPath RL, an open-source Python package available on GitHub. While MCTS has already been implemented for chemical synthesis (Segler, Preuss, and Waller 2018), RetroPath RL is to our knowledge the first application of this algorithm to biochemistry and metabolic engineering. It is a versatile, modular command line tool, build for metabolic engineers, that takes as input a compound of interest, a microbial strain as sink and a set of reaction rules.

While RetroPath RL can take as input any set of chemical rules in encoded into the SMARTS linear notation, giving great freedom to users to use reactions from their pathways or microbial strain of expertise, the tests presented in this article use rules from our RetroRules database (Duigou et al. 2019). RetroRules was built using a data-driven approach showcased in earlier version of RetroPath (Delépine et al. 2018), allowing promiscuity encoding. Using both reaction rules at various diameters and chemical similarity scoring, we can tune the allowed promiscuity within our search, which is an advantage over building reaction rules from EC numbers as is commonly done in metabolic engineering. Another advantage of a data-driven approach is that it is bound to get more precise and information-rich as metabolic databases expand.

We validated RetroPath RL by verifying on a manually curated dataset that a literature-described pathway could be found for 20 different compounds. The fact that the experimental pathways were found for 75% of compounds using default settings and 95% of the time using more tolerant ones, with much better results when using biochemically guided search, confirms the ability of RetroPath RL to suggest experimentally viable pathways for metabolic engineers on numerous compounds.

Moreover, following the standards we set in our previous paper (Delépine et al. 2018), we tested RetroPath RL on a larger dataset and found a pathway for 83.6% of compounds that were results of successful metabolic engineering projects. These results confirm that RetroPath RL generalises well to metabolic engineering compounds outside the manually curated golden dataset. Moreover, RetroPath RL suggests more pathways (median number of 11.5 versus 4.5), giving metabolic engineers more suggestions with which to exercise their expertise.

While a restricted number of microbial strains was provided with RetroPath RL, the user can provide his own strain or supplement the media with a given compound of interest. For example, when supplementing xylene to the microbial strain an experimentally described pathway to TPA (Bramucci, M.G., McCutchen, C.M., Nagarajan, V., Thomas, S.M. 2001) is found, and when adding thebaine to the media, a pathway for morphine production is also found (Thodey, Galanie, and Smolke 2014). A novel feature that should be welcomed by the metabolic engineering community is the ability of RetroPath RL to suggest media supplements to complement the enzymatic synthesis plan. This feature, tested on 8 pathways, found the experimentally described supplement for 5 of them, and also suggests other supplements to test.

We showcase RetroPath RL’s modularity by biasing the search towards less toxic intermediates, and also provide expert users 2 strategies to speed-up the retrosynthetic search, either by storing results in a database (**Supplementary Note 3**) or extending a search from previously run search trees (**Supplementary Note 4**). Another feature of interest is the ability to use RetroPath RL for biosensor design, as we also demonstrated with our previously developed tools (Delépine et al. 2016; Libis, Delépine, and Faulon 2016). This can be used in conjunction with a dataset of detectable compounds (Koch et al. 2018) to allow for design of Sensing Enabling Metabolic Pathways.

Despite the advantages presented above, RetroPath RL still presents some limits, mostly related to pathway ranking. The authors of this article believe modular design, with a downstream analysis of selected pathways, to be a more appropriate course of action than ranking within the tool: so many genome or growth conditions modifications can increase a pathway yield that including all ranking schemes into one to come up with a ‘best’ pathway neglects both metabolic engineers’ expertise and years of lessons from the industry. However, most developers present all integrated tools that perform pathway search and ranking together. Ranking schemes in such tools can involve accounting for enzyme integration into the microbial strain, kinetics, toxicity, carried flux, thermodynamics or preferred cofactor usage (Kumar et al. 2018; Tokic et al. 2018). Our approach provides two advantages over integrated tools. First, ranking the pathways separately allows for a much more modular integration, and the ability to integrate new ideas into ranking much more easily. Moreover, as shown in the custom use section, for advanced users attached to using these schemes, it is possible to integrate them easily given RetroPath RL’s modular code and the ability of the Monte Carlo Tree Search algorithm to bias the search towards or away from properties of interest.

While stereochemistry has been described as one of Nature’s most interesting advantages compared to the traditional chemical industry (Lin, Warden-Rothman, and Voigt 2019), it is not used in results presented here, mainly due to current technical limitations to the way stereochemistry is handled by the cheminformatics packages we used at the moment. However, since our formalism uses rules SMARTS which are stereo-aware (Daylight Chemical Information Systems, Inc. 2008), and our standardisation schemes can leave or remove stereochemistry, users wishing to use stereochemistry in RetroPath RL can do so.

Another major advance that could be included in RetroPath RL would be to guide reaction selection steps through learned values instead of similarity. For example, this was implemented in (Segler, Preuss, and Waller 2018) or (Schreck, Coley, and Bishop 2019). However, the authors learned values from Reaxys (Elsevier Life Sciences n.d.) which contains 12.4 million single-step reactions (compared to around 80k in MetaNetX, including reactions without chemical structures (Moretti et al. 2016)). Therefore, using learned values in bio-retrosynthesis seems out of reach for the moment, but could become available in the coming years due to the intense curation efforts under way in the community. Moreover, those learned values in bio-retrosynthesis would have to be strain-dependent, as an intermediate compound’s value for bio-retrosynthesis depends highly on the strain of interest.

In conclusion, we present here a highly modular tool using one of the latest tree search algorithms for bio-retrosynthesis. This tool is modular enough for expert users to input their expert knowledge and has been thoroughly tested on datasets of interest to the community.

## Materials and methods

All chemical operations were performed using RDKit release 2019.03.1.0 and Python 3.6.

### Compound standardisation

All compounds were standardised using the following steps:

- sanitising chemical depictions using RDKit’s SanitizeMol method
- removing isotope
- neutralising charges
- removing stereo
- converting back and forth to InChI (Heller et al. 2013) to ensure tautomerism consistency

### Reaction rule encoding

Our reaction rules are generated as presented in our RetroRules database (Duigou et al. 2019). Briefly, we extracted known biochemical reactions from the MetaNetX (Moretti et al. 2016) database version 3.1 and filtered out incomplete reactions. We identified the reaction centres based on atom-atom mappings we performed using the Reaction Decoder software ((Rahman et al. 2016), version 2.1.0). We decomposed multi-substrates reactions into mono-substrate components by considering one substrate per component and only the subset of products that share at least one atom with the substrate. Mono-components involving a typical cofactor (such as CO2, ATP, NADH...) as substrate were excluded. Finally, each mono-component reaction is encoded into a collection of reaction rules using the SMARTS (Daylight Chemical Information Systems, Inc. 2008) formalism for a diameter ranging from 2 to 16 around the reaction centre.

The main difference with the RetroRules procedure aforementioned is that we use implicit hydrogen notation in the reaction rules instead of explicit, which allows a much faster computation of standardisation of compounds and reaction rule application through RDKit.

To validate the reaction rules, we checked that applying the reaction rules on the substrates used as templates produce the products described in the template reactions. The success rates range from 99.4% for reaction rules at diameter 2 to 99.8% for those at diameter 16. We also performed a cross-radius consistency check by ensuring that sets of products produced by large diameter reaction rules (e.g. reaction rules generated with a diameter of 10) are always subsets of products produced by reaction rules at any smaller diameter (e.g. reaction rules for diameter 8), which show a success rate of 99.3%. Only reaction rules passing successfully both procedures have been retained and used, which represent almost 146k distinct reaction rules (available for download on RetroRules website, download page, release rr02) and model more than 18k biochemical reactions in both directions.

### Branching factor calculation

We generated what we called the Extended Metabolic Space (EMS) at 1 step, i.e. the metabolic space that can be reached by applying our reaction rules once on all compounds of MetaNetX (Moretti et al. 2016) being used as a template for generating at least one reaction rule. We filtered out substrates having a molecular weight greater than 1 kDa. We set a timeout cut-off of 1 second on rule application. We then analysed the results presented in Table 1 and **Supplementary Table 1** using the NetworkX python library (Aric A. Hagberg 2008).

### Chemical score calculation

After compound standardisation, we calculate a 1024 binary Morgan fingerprints vector using the RDKit method GetMorganFingerprintAsBitVect at diameter 4.

For substrate score, the query substrate is compared to all native substrates a rule was learned on using Tanimoto score (Maggiora and Shanmugasundaram 2004; Bajusz, Rácz, and Héberger 2015), and the maximum score is kept.

For products and substrate scoring, the procedure is as follows:

For each native (n_sub, n_products) couple:

- calculate Tanimoto of query and native substrate.
- generate all combinations of native and query products (below 1000 to avoid a combinatorial explosion. The number of combinations is *n!* where *n* is the number of products generated by the rule application)
- for each combination, calculate the geometric mean of the Tanimoto scores of products
- Keep the highest combination score.
- The score of this native combination is the product of the substrate score and the highest combination score.

The score of the rule is the highest score of all native substrates and products the rule was generated from.

Rules generating a different number of products from the template receive a score of −1.

### Biological score calculation

A penalty is calculated as presented in our RetroPath 2.0 paper (Delépine et al. 2018). Briefly, we clustered reaction rules according to the EC number annotations inherited from the template reactions. Independently, we clustered enzyme sequences collected from UniProt (The UniProt Consortium 2017) (release 2019_04) according to sequence similarity using the cd-hit software (version 4.6.8)(Li and Godzik 2006). We establish the sequences to reaction rules associations based on Rhea (v98)(Morgat et al. 2017), MetaCyc (v21.5)(Caspi et al. 2018) and Reactome (v66)(Fabregat et al. 2018) crosslinks. The penalty score was then computed as

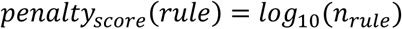

with *n*_*rule*_ the number of distinct clusters that contains sequences associated to the rule. In addition, we normalised this penalty into a score comprised between 0 and 1 using the following function:

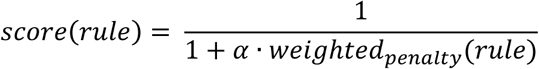

with

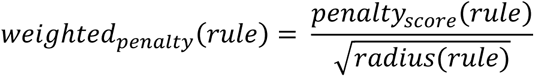

and

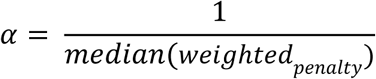

### Sinks construction

Except stated, sink compounds have been extracted from genome-scale metabolic models by only collecting the chemicals that lie in the cytosol compartment. In addition, we filtered out “dead-end” compounds, i.e. compounds that cannot be produced by any reactions in a steady-state metabolic model, that we detected by performing a Flux Variability Analysis using the COBRApy package (v0.15.3) (Ebrahim et al. 2013).

Chemical structures have been obtained using crosslinks from the models to metabolic databases. In case no crosslink or not any valid structure was found, the PubChem (Kim et al. 2019) database was examined using compound names as query. Finally, all chemical structures were standardised as described in the “Compound standardisation” section.

### Available sinks

The available sinks provided with our software are iML1515 (Monk et al. 2017), iJO1366 (Orth et al. 2011) and core *E. coli* metabolism (Orth, Fleming, and Palsson 2010), as well as *Bacillus Subtilis* iYO844 model (Oh et al. 2007) and our set of detectable compounds for biosensor design (Koch et al. 2018). The genome scale models were obtained from the BiGG Models database (King et al. 2016). By default, we used sinks from the iML1515 model.

### RetroPath 2.0 configuration

For all tests made with RetroPath 2.0, we perform all executions on a recent work station. We use the sink extracted from the iML1515 model, and we set the maximum pathway length to 5, the maximum number of structures to keep for next iteration to 1000, and a 3 hours per execution time budget. We then use the rp2paths software available on GitHub at https://github.com/brsynth/rp2paths to extract the pathways from RetroPath 2.0 output (Delépine et al. 2018).

### Monte Carlo Tree Search implementation

The aim of Markov decision processes is to model sequential decision processes of an agent in an environment (Sutton et al. 1998). Its most notable components are states (representing positions in a game) and actions (allowed transformations from the state).

In RetroPath RL, following the method developed by (Segler, Preuss, and Waller 2018), we consider states to be a set of molecules. The initial state contains only the target compound one desires to produce. Actions are transformations of any molecule of the state that do not transform the compound into itself nor produce a non-sink compound that has already been produced before in the synthesis plan so as to avoid loop searches.

A state is considered terminal if all compounds are in the sink, if no move can be applied to this state or if the maximum allowed depth has been reached.

Monte Carlo Tree Search is a reinforcement learning approach that builds a search tree and stochastically explores search space to bias search towards most promising regions of combinatorial space, following steps presented in Figure 2B and detailed below.

### Selection

Starting from the root node, the best child nodes according to the selection policy are iteratively chosen until a leaf node is reached. The formula we used is:

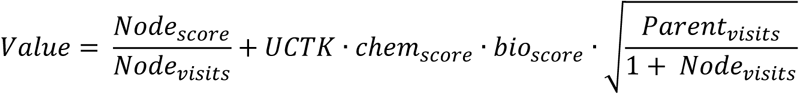

where *Node*_*score*_ is the cumulative score from rollouts, *Node*_*visits*_ is the number of visits to this node, *UCTK* is the UCT (Upper Confidence Trees) constant used (balances between exploitation and exploration), chemical and biological scores are the scores of the move leading to this node and *parent*_*visits*_ is the number of visits of this Node’s parents.

Other policies have been developed, notably with only chemical, biological score, or no scoring. In our implementation, grandchildren of a node can only be explored if all his children have had at least *minimal_visits* number of visits. This allows mandatory rollout on different branches at least once to favour exploration.

### Expansion

For each compound that is not in the sink, its *n* best moves are identified and stored (with *n* the maximal number of children allowed for the node). Then, the *n* best moves overall on the state are selected and children created iteratively (one at the first visit of the node, the next at the next visit and so on) for each of these moves and a rollout is performed.

### Rollout

Rollout is an iterative procedure that starts by checking the status of the state. If it is terminal, a reward (or penalty) is returned according to the rewarding policy. If it is not terminal, a transformation is randomly sampled from available transformations and the process is repeated. This is performed until a maximum number of rollout steps or the maximal depth of the tree is reached. The function for random sampling used throughout this study gives weight chemical * biological score to moves, therefore giving more probability of being sampled to higher scoring moves according to our scoring scheme.

### Rewarding policy

A state is rewarded as follows:

1. receives a penalty of −1 when no compound is solved
2. receives a bonus of 5 when the state is fully solved.
3. receives a score of number_found/total_number when only a fraction of molecules in the state are solved.

### Update

The node and its parents update their value and node counts according to the results obtained from the above rewarding scheme after rollout.

### Returning complete pathways

Each time a full pathway is found during the tree expansion, the pathway is returned, and an additional bonus of 10 is received by the node, to allow for biasing towards similar successful pathways. At the end of the search, the most visited pathway is returned (“best”), and all pathways are returned ranked in order of decreasing biochemical score.

### Golden dataset construction

In order to perform golden dataset curation, we focused on articles where pathways were explicitly described (i.e. no missing steps, and available intermediate compounds). We retrieved compound structures in PubChem (Kim et al. 2019) and EC numbers from Brenda (Jeske et al. 2019) based on enzyme name as given in the article. We selected pathways of strictly more than 1 step. We then verified for each step that we had chemical rules available in our RetroRules (Duigou et al. 2019) database to encode the described transformations. This ensures a fair comparison between tools using our publicly available reaction rules, in order to evaluate separately retrosynthesis tools and the underlying chemical rules. The detailed list with references is available as **Supplementary Table 4**, and the pathways as **Supplementary Data 1**.

### Experimental pathway comparison

In order to compare the pathways found by RetroPath 2.0 or RL and the experimental pathways described in the literature, we have two types of information: compound identity and EC number identity. We consider the reaction EC number to be equal if it is identical up to 3 digits (1.1.1.x is identical to 1.1.1.y). Since spontaneous reactions do not have an EC number, we used compound identity for comparison, and EC number as additional information.

### LASER retrieval and Metabolic Engineering completion

We build the LASER dataset by parsing target molecule and microbial strain information from the LASER database published by Winkler et al. which provides a curated list of more than 600 successfully implemented metabolic designs (Winkler, Halweg-Edwards, and Gill 2016). When available, we store the chemical structure provided by MetaCyc (v2RL), otherwise we query the PubChem database based on target compound names. We augmented this list with target compounds reported in the Metabolic Engineering journal in 2016 (volumes 33 to 38) and published in RetroPath 2.0 (Delépine et al. 2018). All chemical structures have been standardised using the procedure described in the “Compound standardisation” section. The final dataset used in the present paper contains 211 unique structures that are provided as **Supplementary Data 2**.

### Extracting supplements from LASER

LASER provides a ‘Media’ line that contains addition to the media, extracted using Natural Language Processing. This can include antibiotics, promoter inducers or supplements of interest required to build the pathway. We removed all compounds that did not concern pathway supplements and removed early precursor supplementation. Structures were obtained from PubChem. We obtain a list of only 6 pathways satisfying these requirements, and 8 when we also add two pathways for TPA from literature (Bramucci, M.G., McCutchen, C.M., Nagarajan, V., Thomas, S.M. 2001; J., J., and J. 2006). This list is available as **Supplementary Data 4**.

### Supplement finder feature

The Supplement finder functions as follows:

1. load search tree in memory
2. explore all nodes and keep compound structures that complete of a chemical state (i.e. all other compounds of the state are solved)
3. compounds that allow for completion of more than N states (which would complement N pathways) are kept. Here N= 1: any compound that can complement a pathway is kept.
4. compounds are filtered according to presence in a Database of interest. Here, we filtered according to presence in the Sigma catalogue.
5. we keep the N best suggestions (according to number of pathways that are completed). Here, we returned up to 20 potential supplements.
6. all completed pathways are extracted for future analysis.

### Toxicity implementation

We used data from EcoliTox (Planson et al. 2012) and XTMS (Carbonell et al. 2014) to build a QSAR model (using as input features 1024 binary Morgan fingerprints vector calculated with the RDKit method GetMorganFingerprintAsBitVect at diameter 4), predicting log(IC50) of the compounds. We train our model using scikit-learn (version 0.19.1) (Pedregosa et al. 2011). The model is a multi-layer perceptron trained with the default parameters from scitkit-learn except the following parameters: maximum iteration of 20000, adaptive learning rate, adam solver, early stopping and the following layers: (10, 100,100, 20). This model has a Leave-One-Out (LOO) score of 0.81. In prediction mode, toxicity was used only if the predicted log(IC50) <= 0. The modified UCT function that was used is:

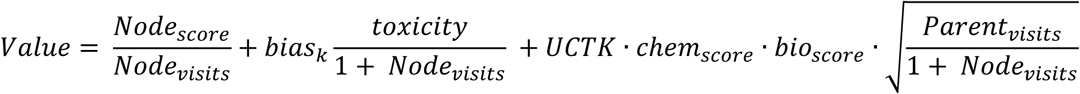

### Hardware

We ran our tests on 2 calculation clusters and 1 personal computer. Tests involving using the NoSQL Database and media supplementation were run on a desktop computer with the following characteristics: CPU is Intel(R) Xeon(R) W-2145 CPU @ 3.7 GHz and 32G of RAM. Tests for the golden dataset evaluation were performed on the Migale (https://migale.inra.fr) cluster (2.0 to 2.4 GHz CPUs). Allocated resources per job were set to 1 vCPU and 20 GB. Tests for the LASER database evaluation were run on the IFB cluster (http://taskforce-nncr.gitlab.cluster.france-bioinformatique.fr/doc/cluster-desc/) (2.3 GHz CPUs). Allocated resources per job were set to 1 vCPU and 40 GB. Cluster tests were run using snakemake 5.4.0 (Köster and Rahmann 2012).

## Supporting information

Golden dataset pathways under json and SBML format

LASER database and Metabolic Engineering compounds

LASER database and Metabolic Engineering results

Targets and supplements for experimentally described pathways

## Code and data availability

RetroPath RL is available at https://github.com/brsynth/RetroPathRL under a MIT license. Reaction rules are available at https://retrorules.org/dl.

## Supporting Information

The Supporting Information for this article contains:

- **Supplementary Table 1**: Mean number of rules applying to a compound at various diameters.
- **Supplementary Table 2**: Detailed configuration data for RetroPath RL runs on validation datasets.
- **Supplementary Table 3**: Toxicity biased results.
- **Supplementary Table 4**: Golden dataset structures and references.
- **Supplementary Note 1** (with **Supplementary Figures 1 to 12)**: Detailed parameters explanation and analysis.
- **Supplementary Note 2**: Detailed golden dataset analysis
- **Supplementary Note 3** (with **Supplementary Figure 13**): Database sped-up calculations
- **Supplementary Note 4** (with **Supplementary Figure 14**): Extending a previous search
- **Supplementary Data 1**: Golden dataset pathways under json and SBML format.
- **Supplementary Data 2**: LASER database and Metabolic Engineering compounds.
- **Supplementary Data 3**: LASER database and Metabolic Engineering results.
- **Supplementary Data 4**: Targets and supplements for experimentally described pathways.

## Author contribution

M.K., T.D. and J.-L. F. designed the study. M.K. developed the MCTS algorithm. T.D. developed rule extraction and application code, strain analysis and LASER extraction. Both M.K. and T.D. tested and validated the software. M.K., T.D. and J.-L. F. wrote the paper.

## Acknowledgements

We are grateful to the INRA MIGALE bioinformatics platform (https://migale.inra.fr) for providing computational resources. We would like to thank the French Institute of Bioinformatics (IFB, ANR-11-INBS-0013, https://www.france-bioinformatique.fr) for providing storage and computing resources on its Core Cluster. M.K is supported by DGA (French Ministry of Defense) and Ecole Polytechnique. J.-L.F. acknowledges support from BBSRC/ EPSRC (grant number BB/M017702/1) and from the ANR (grant numbers ANR-15-CE21-0008 and ANR-17-CE07-0046). We thank Pablo Carbonell for providing the toxicity data.

