## Supplementary Information for "Reinforcement Learning for Bio-Retrosynthesis"

### Reinforcement Learning for Bio-Retrosynthesis - Supplementary Information

The Supplementary Data for this article contains:

- **Supplementary Table 1**: Mean number of rules applying to a compound at various diameters.
- **Supplementary Table 2**: Detailed configuration data for RetroPath RL runs on validation datasets.
- **Supplementary Table 3**: Toxicity biased results.
- **Supplementary Table 4**: Golden dataset structures and references.
- **Supplementary Note 1** (with **Supplementary Figures 1 to 12)**: Detailed parameters explanation and analysis.
- **Supplementary Note 2**: Detailed golden dataset analysis
- **Supplementary Note 3** (with **Supplementary Figure 13**): Database sped-up calculations
- 
- **Supplementary Note 4** (with **Supplementary Figure 14**): Extending a previous search
- **Supplementary Data 1**: Golden dataset pathways under json and SBML format.
- **Supplementary Data 2**: Laser database and Metabolic Engineering compounds.
- **Supplementary Data 3**: Laser database and Metabolic Engineering results.
- **Supplementary Data 4**: Targets and supplements for experimentally described pathways.

**Supplementary Table 1:** Average number of rules that apply to a compound according to the diameter at which the rule is used. This shows that more promiscuous rules (at lower diameters) generate a bigger combinatorial space.

| **Diameter used** | 2 | 4 | 6 | 8 | 10 | 12 | 14 | 16 |
|---|---|---|---|---|---|---|---|---|
| **Average number of rules that apply** | 797 | 227 | 73 | 35 | 20 | 14 | 11 | 8 |

**Supplementary Table 2:** Detailed configuration data for RetroPath RL runs on validation datasets.

| **Parameter name** | **Golden Default** | **Golden Rescue** | **Laser Default** | **Laser Rescue** |
|---|---|---|---|---|
| Itermax | 10000 | 10000 | 50000 | 10000 |
| Expansion width | 10 | 15 | 10 | 10 |
| Time budget (s) | 10800 when comparing to RP2, 14400 otherwise | 10800 when comparing to RP2, 14400 otherwise | 10800 | 10800 |
| Max depth | 7 | 7 | 10 | 7 |
| UCT Policy | Biochemical | Biochemical | Biochemical | Biochemical |
| UCTK | 2 | 2 | 2 | 2 |
| Rollout Policy | Uniform on biochemical score | Uniform on biochemical score | Uniform on biochemical score | Uniform on biochemical score |
| Max rollout | 2 | 2 | 2 | 2 |
| Chemical scoring | Substrate and product | Substrate and product | Substrate and product | Substrate and product |
| Virtual visits | 0 | 0 | 0 | 0 |
| Rule diameters | 6, 10, 16 | 6, 10, 16 | 6, 10, 16 | 2, 6, 10, 16 |
| Biological score cut-off | 0.3 | 0.15 | 0.3 | 0 |
| Substrate score cut-off | 0.3 | 0.15 | 0.3 | 0 |
| Chemical score cut-off | 0.3 | 0.15 | 0.3 | 0 |
| Minimal visits count | 1 | 1 | 1 | 1 |
| Fire timeout (s) | 1 | 1 | 1 | 1 |
| Penalty | -1 | -1 | -1 | -1 |
| Reward | 5 | 5 | 5 | 5 |
| Full pathway reward | 10 | 10 | 10 | 10 |
| Seed | 42 | 42 | 42 | 42 |

**Supplementary Table 3: Toxicity biased results.** Comparing results between the default configuration and the one using the toxicity score to bias the search. Total pathway number is the total number of pathways found with the given time and iteration budget. The iteration and rank for the experimental pathway refer to the pathway described in the golden dataset.

| **Compound** | **Total pathway number.** default - *toxicity* | **Iteration for finding the experimental pathway.** default - *toxicity* | **Rank of experimental pathway.** default - *toxicity* |
|---|---|---|---|
| 3-methyl-1-butanol | 1 - *1* | 4 - *4* | 1 - *1* |
| 1,4-Butanediol | NA - *NA* | NA - *NA* | NA - *NA* |
| 2-amino-1,3-propanediol | 1 - *1* | 1 - *1* | 1 - *1* |
| 2,5-DHBA | 34 - *31* | 88 - *97* | 4 - *4* |
| benzyl alcohol | 13 - *22* | 1815 - *2213* | 4 - *8* |
| caroten | 4 - *4* | 6508 - *6443* | 1 - *1* |
| cis,cis muconate | 11 - *11* | 159 - *176* | 6 - *6* |
| glutaric acid | 61 - *74* | 1114 - *571* | 13 - *7* |
| lycopene | 106 - *103* | 101 - *96* | 1 - *1* |
| mesaconic acid | 14 - *14* | 9 - *9* | 1 - *1* |
| naringenin | 38 - *40* | 408 - *1011* | 1 - *9* |
| N-methylpyrrolinium | 9 - *10* | NA - *NA* | NA - *NA* |
| p-hydroxystyrene | 59 - *61* | 34 - *33* | 1 - *1* |
| piceatannol | 34 - *38* | 8581 - *8996* | 32 - *36* |
| pinocembrin | 25 - *25* | 66 - *55* | 2 - *2* |
| protopanaxadiol | 1 - *2* | NA - *NA* | NA - *NA* |
| styrene | 43 - *45* | 14 - *13* | 2 - *2* |
| TPA | 9 - *10* | NA - *NA* | NA - *NA* |
| vanillin | 34 - *36* | 169 - *171* | 4 - *4* |
| violacein | 3 - *3* | 158 - *158* | 3 - *3* |

**Supplementary Table 4:** Golden dataset. This dataset contains the compounds, structures and references used for experimental pathway analysis presented in Results - golden set.

| Name | InChI | Reference |
|---|---|---|
| 3-methyl-1-butanol | InChI=1S/C5H12O/c1-5(2)3-4-6/h5-6H,3-4H2,1-2H3 | (Connor and Liao 2008) |
| 1,4-Butanediol | InChI=1S/C4H10O2/c5-3-1-2-4-6/h5-6H,1-4H2 | (Hwang et al. 2014; Yim et al. 2011) |
| 2-amino-1,3-propanediol | InChI=1S/C3H9NO2/c4-3(1-5)2-6/h3,5-6H,1-2,4H2 | (Luo et al. 2019) |
| 2,5-DHBA | InChI=1S/C7H6O4/c8-4-1-2-6(9)5(3-4)7(10)11/h1-3,8-9H,(H,10,11) | (Shen et al. 2018) |
| benzyl alcohol | InChI=1S/C7H8O/c8-6-7-4-2-1-3-5-7/h1-5,8H,6H2 | (Pugh et al. 2015) |
| caroten | InChI=1S/C40H56/c1-31(19-13-21-33(3)25-27-37-35(5)23-15-29-39(37,7)8)17-11-12-18-32(2)20-14-22-34(4)26-28-38-36(6)24-16-30-40(38,9)10/h11-14,17-22,25-28H,15-16,23-24,29-30H2,1-10H3/b12-11+,19-13+,20-14+,27-25+,28-26+,31-17+,32-18+,33-21+,34-22+ | (Yang and Guo 2014) |
| cis,cis muconate | InChI=1S/C6H6O4/c7-5(8)3-1-2-4-6(9)10/h1-4H,(H,7,8)(H,9,10)/p-2/b3-1-,4-2- | (Lin et al. 2014) |
| glutaric acid | InChI=1S/C5H8O4/c6-4(7)2-1-3-5(8)9/h1-3H2,(H,6,7)(H,8,9) | (Park et al. 2013) |
| lycopene | InChI=1S/C40H56/c1-33(2)19-13-23-37(7)27-17-31-39(9)29-15-25-35(5)21-11-12-22-36(6)26-16-30-40(10)32-18-28-38(8)24-14-20-34(3)4/h11-12,15-22,25-32H,13-14,23-24H2,1-10H3/b12-11+,25-15+,26-16+,31-17+,32-18+,35-21+,36-22+,37-27+,38-28+,39-29+,40-30+ | (Ma et al. 2019) |
| mesaconic acid | InChI=1S/C5H6O4/c1-3(5(8)9)2-4(6)7/h2H,1H3,(H,6,7)(H,8,9)/b3-2+ | (Wang and Zhang 2015) |
| naringenin | InChI=1S/C15H12O5/c16-9-3-1-8(2-4-9)13-7-12(19)15-11(18)5-10(17)6-14(15)20-13/h1-6,13,16-18H,7H2 | (Santos, Koffas, and Stephanopoulos 2011) |
| N-methylpyrrolinium | InChI=1S/C5H10N/c1-6-4-2-3-5-6/h4H,2-3,5H2,1H3/q+1 | (Ping et al. 2019) |
| p-hydroxystyrene | InChI=1S/C8H8O/c1-2-7-3-5-8(9)6-4-7/h2-6,9H,1H2 | (Qi et al. 2007) |

| | | |
|---|---|---|
| piceatannol | InChI=1S/C14H12O4/c15-11-5-10(6-12(16)8-11)2-1-9-3-4-13(17)14(18)7-9/h1-8,15-18H/b2-1+ | (Shrestha, Pandey, and Sohng 2019) |
| pinocembrin | InChI=1S/C15H12O4/c16-10-6-11(17)15-12(18)8-13(19-14(15)7-10)9-4-2-1-3-5-9/h1-7,13,16-17H,8H2/t13-/m0/s1 | (Kim, Lee, and Ahn 2014) |
| protopanaxadiol | InChI=1S/C30H52O3/c1-19(2)10-9-14-30(8,33)20-11-16-29(7)25(20)21(31)18-23-27(5)15-13-24(32)26(3,4)22(27)12-17-28(23,29)6/h10,20-25,31-33H,9,11-18H2,1-8H3/t20-,21+,22-,23+,24-,25-,27-,28+,29+,30+/m0/s1 | (Wu et al. 2019) |
| styrene | InChI=1S/C8H8/c1-2-8-6-4-3-5-7-8/h2-7H,1H2 | (McKenna and Nielsen 2011) |
| TPA | InChI=1S/C8H6O4/c9-7(10)5-1-2-6(4-3-5)8(11)12/h1-4H,(H,9,10)(H,11,12) | (Bramucci, M.G., McCutchen, C.M., Nagarajan, V., Thomas, S.M. 2001) |
| vanillin | InChI=1S/C8H8O3/c1-11-8-4-6(5-9)2-3-7(8)10/h2-5,10H,1H3 | (Kunjapur, Tarasova, and Prather 2014) |
| violacein | InChI=1S/C20H13N3O3/c24-10-5-6-15-12(7-10)14(9-21-15)17-8-13(19(25)23-17)18-11-3-1-2-4-16(11)22-20(18)26/h1-9,21,24H,(H,22,26)(H,23,25)/b18-13+ | (Jones et al. 2015; Hoshino 2011) |

### Supplementary Note 1 : Parameter Role and Effects

The aim of this supplementary note is to detail the different parameters available in RP3 and their roles and effects.

A number of methods and ideas were taken or inspired from the following master thesis, which uses Monte Carlo Tree Search against a computer game(Kozelek 2009).

#### Rule selection parameters

Biological score

As described in the main text, this score characterises our confidence that a sequence exists to catalyse the reaction of interest. It is normalised between 0 and 1. Using a cut-off on this score removes less trustworthy reaction rules. We see in **Supplementary Figure 1** that results do not vary between using a cut-off from 0 to 0.3, and we start losing pathways of interest when the cut-off is superior or equal to 0.5. A cut-off of 0.3 therefore seems to be a good trade-off between confidence in the existence of a sequence and obtaining enough retrosynthesis results.

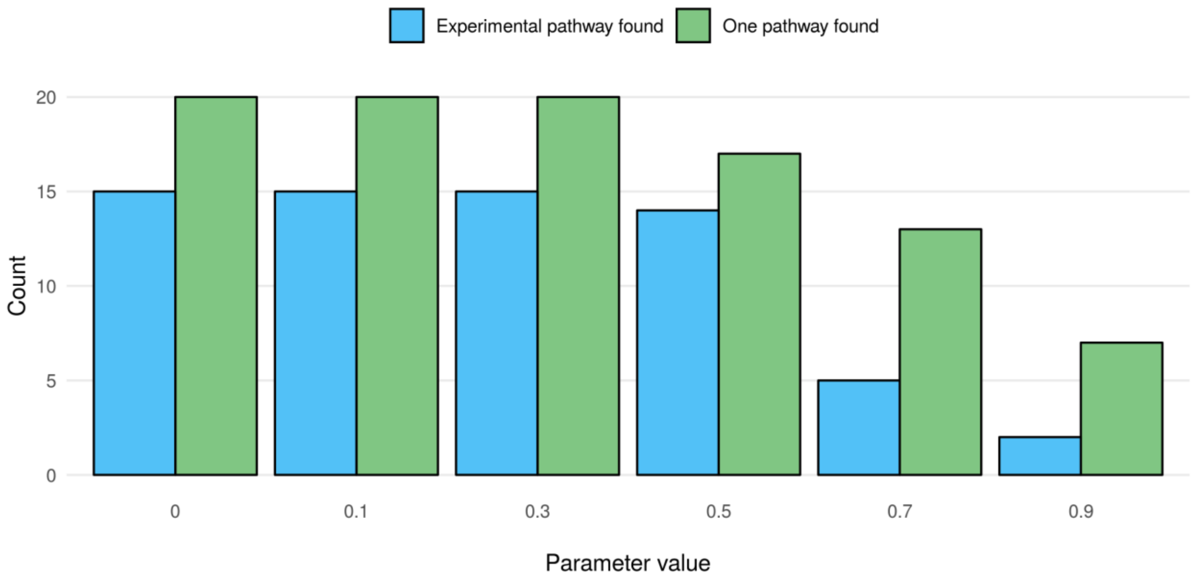


**Supplementary Figure 1: Impact of biological score cut-off on retrieval performance of RetroPath RL**. We compared results between using a biological score cut-off varying between 0 and 0.9. One pathway found means that at least one pathway has been predicted. Experimental pathway found means that the experimental pathway is from amongst the predicted pathways. Also presented as **Figure 5A**.

Chemical score

As described in the main text, this score characterizes our confidence that the reaction rule learned on a substrate and product from a database of interest can truly be applied to a new substrate, based on similarity between substrates and products of the native reaction versus the query reaction. We can see in **Supplementary Figure 2** that allowing reactions that are too different leads the tree to explore too diverse pathways, while being too conservative leads to loss of useful reactions. 0.3 therefore seems to be a good cut-off between confidence that the reaction rule does apply to the compound and allowing exploration of the metabolic space.

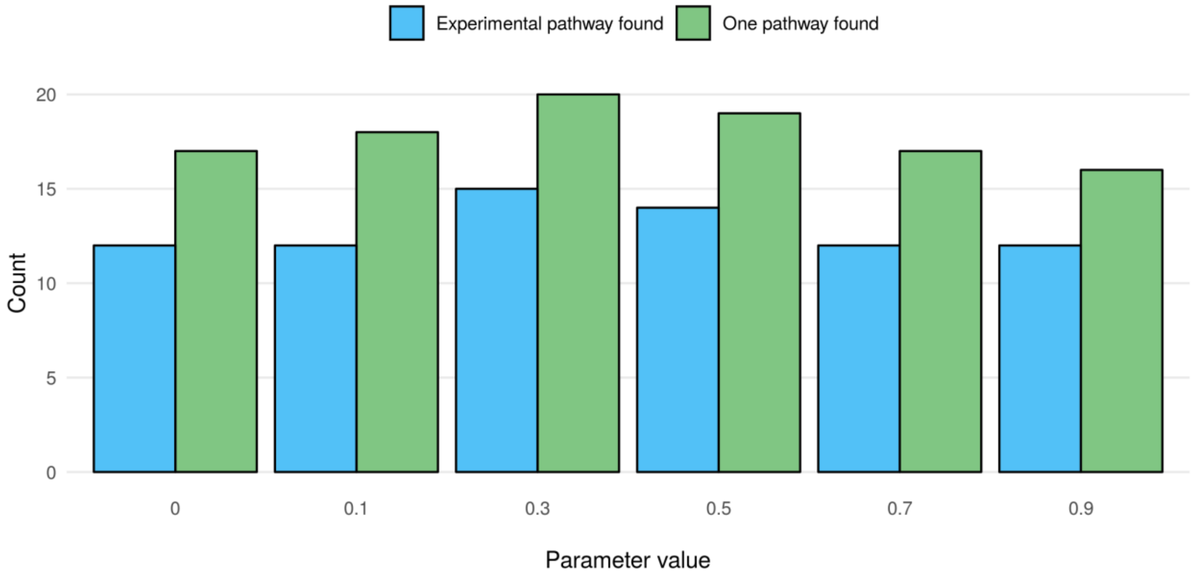


**Supplementary Figure 2: Impact of chemical score cut-off on retrieval performance of RetroPath RL**. We compared results between using a chemical score cut-off varying between 0 and 0.9. One pathway found means that at least one pathway has been predicted. Experimental pathway found means that the experimental pathway is from amongst the predicted pathways. Also presented as **Figure 5B**.

Biological and chemical score

Here we varied both the chemical and biological scores, set at the same value. We can see in **Supplementary Figure 3**, as in Supplementary Figures 1 and 2, that cut-offs of 0.3 provide the best trade-off between exploration and confidence.

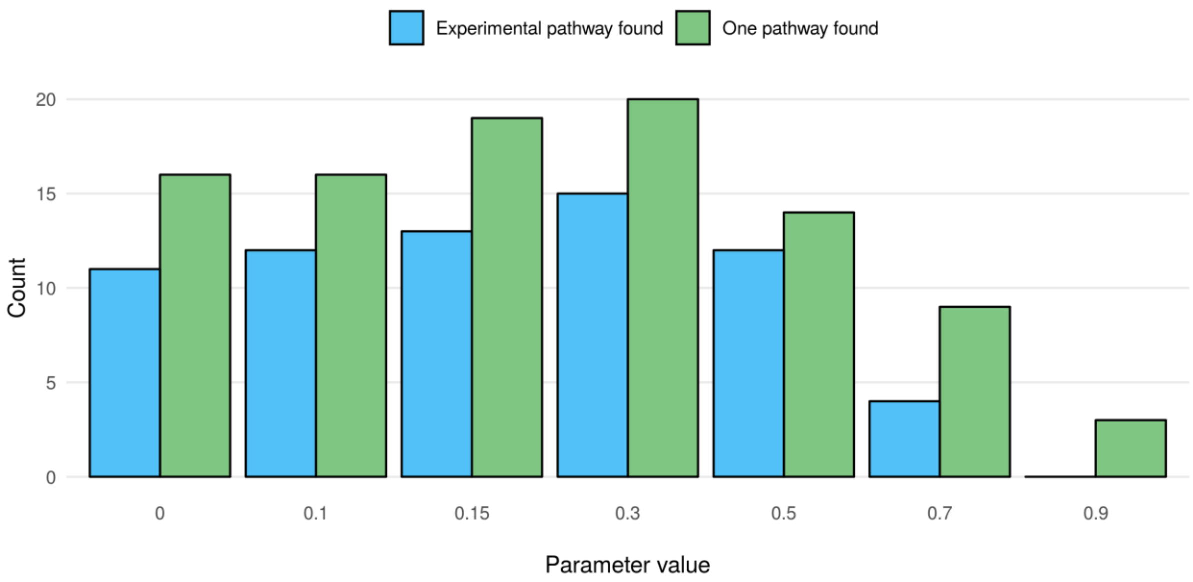


**Supplementary Figure 3: Impact of biochemical score cut-off on retrieval performance of RetroPath RL**. We compared results between using chemical score cut-off and biological score cut-off (set at the same value) varying between 0 and 0.9. One pathway found means that at least one pathway has been predicted. Experimental pathway found means that the experimental pathway is from amongst the predicted pathways. Also presented as **Figure 5C**.

UCT policy

While these policies are used to tune the exploration/exploitation balance, we modified it to guide the search and see the importance of that guidance on finding results. We can see in **Figure 5D** and **Supplementary Figure 4** the best UCT policy to guide our exploration of the metabolic space is our formula including the biochemical score.

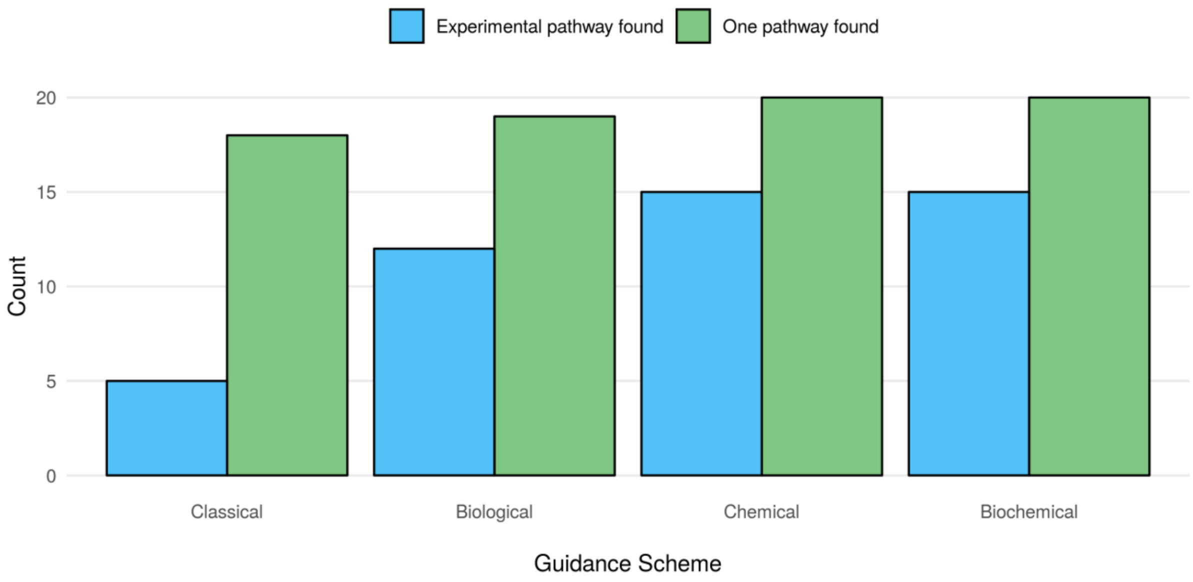


**Supplementary Figure 4: Impact of guiding scheme on retrieval performance of RetroPath RL.** We compared results between guiding the search based on the Classical UCT formula, a formula guided by Biological scoring, Chemical scoring or Biochemical scoring. One pathway found means that at least one pathway has been predicted. Experimental pathway found means that the experimental pathway is from amongst the predicted pathways. Also presented as **Figure 5D**.

Diameters

Diameters characterize the degree of promiscuity we allow in a reaction rule: higher diameters are more specific, while lower diameters are more promiscuous. We found a good trade-off was to allow rules at different levels of promiscuity, using diameters of 6, 10 and 16 (low, medium and high specificity), as shown in **Supplementary Figure 5**.

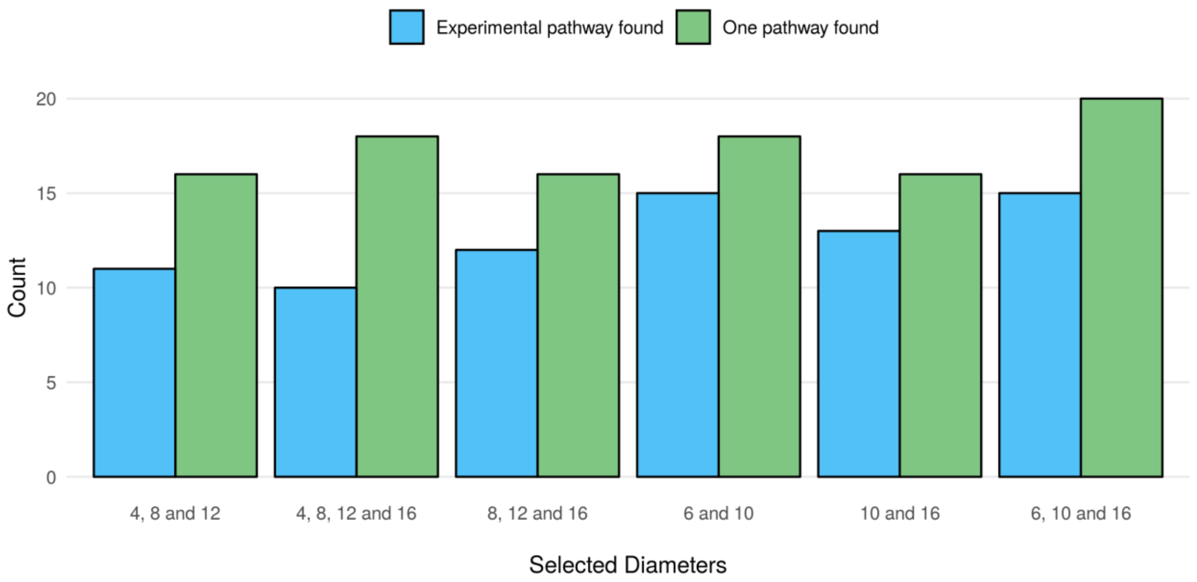


**Supplementary Figure 5: Impact of allowed rule diameters on retrieval performance of RetroPath RL.** We compared results between using different diameter sets. One pathway found means that at least one pathway has been predicted. Experimental pathway found means that the experimental pathway is from amongst the predicted pathways.

#### Exploration parameter

Expansion width

It is the number of children a node is allowed to have. We found 10 and 15 provided a good trade-off between exploration and exploitation, as shown in **Supplementary Figure 6**. We usually tested with 10 children and expanded to 15 for failed compounds.

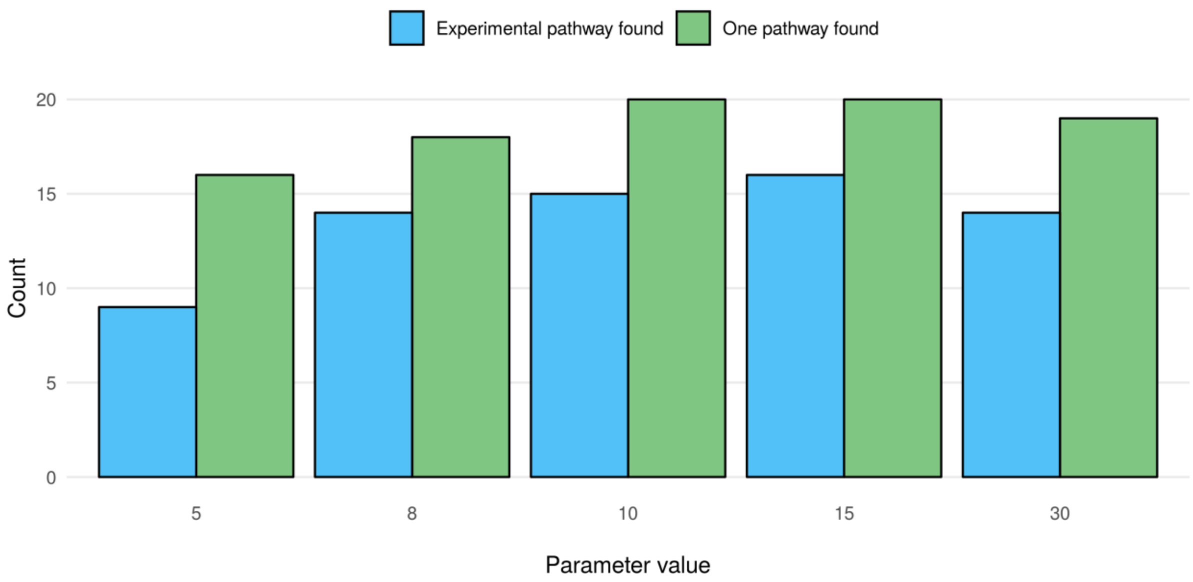


**Supplementary Figure 6: Impact of expansion width on retrieval performance of RetroPath RL.** We compared results between using different expansion width (number of allowed children per node). One pathway found means that at least one pathway has been predicted. Experimental pathway found means that the experimental pathway is from amongst the predicted pathways.

<u>Minimal visit counts</u>

In our implementation, grand-children of a node can only be explored if all his children have had at least minimal_visits visits, where this value was set at 1 in the default settings. This allows mandatory rollout on different branches at least once to favour exploration. We can see from **Supplementary Figure 7** that results are similar when not forcing this exploration with a parameter set to 0.

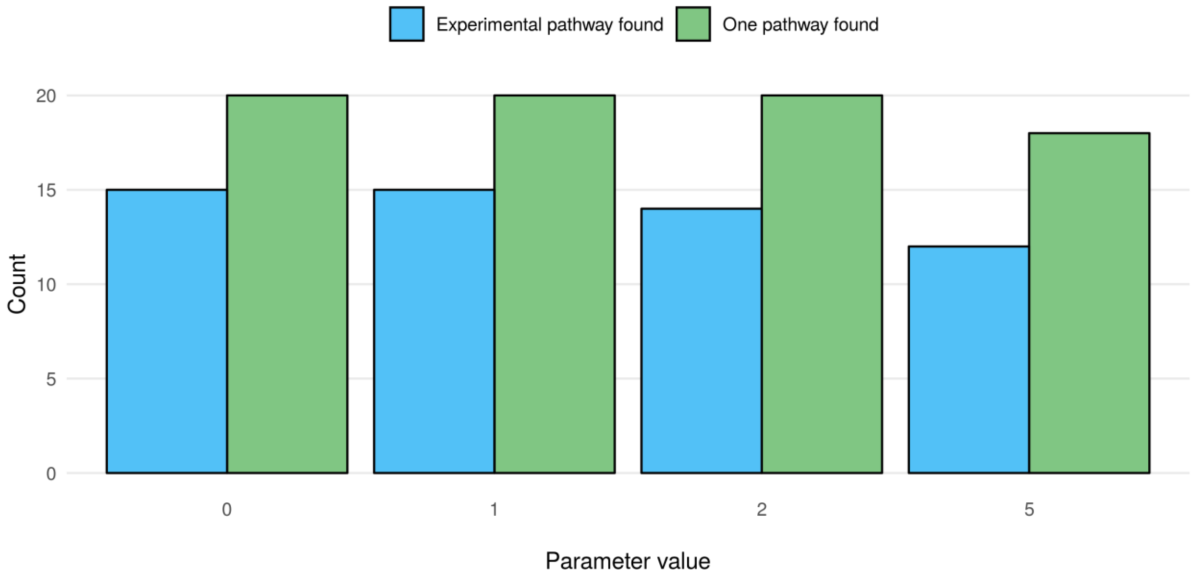


**Supplementary Figure 7: Impact of minimal visit counts on retrieval performance of RetroPath RL.** We compared results between using different minimal visit count values. One pathway found means that at least one pathway has been predicted. Experimental pathway found means that the experimental pathway is from amongst the predicted pathways.

Rollout

This is the rollout depth: the number of reactions performed before analysing the state and returning the state's reward or penalty. **Supplementary Figure 8** shows that rollout depth does not impact our capacity of finding experimental results on the golden dataset. However, unshown results (considering the iteration at which those results are found) suggest best rollout values are either 2 or 3.

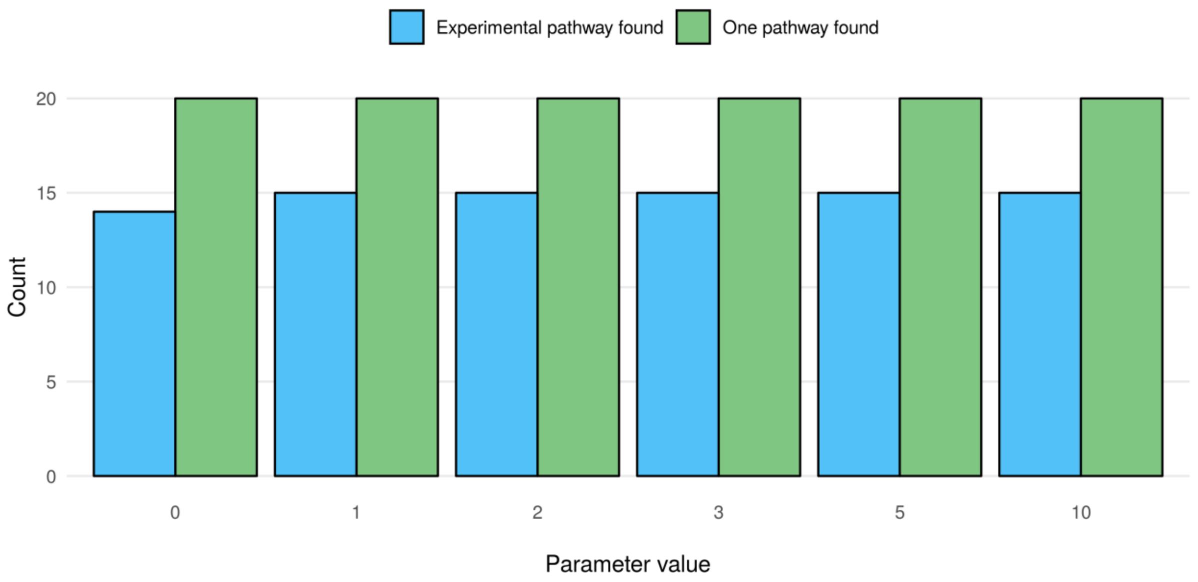


**Supplementary Figure 8: Impact of rollout depth on retrieval performance of RetroPath RL.** We compared results between using different rollout depths. One pathway found means that at least one pathway has been predicted. Experimental pathway found means that the experimental pathway is from amongst the predicted pathways.

UCTK

This constant balances the trade-off between exploration and exploitation in the UCT formula. We can see in **Supplementary Figure 9** that the value allowing the best retrieval rate from the golden dataset is a constant of 2.

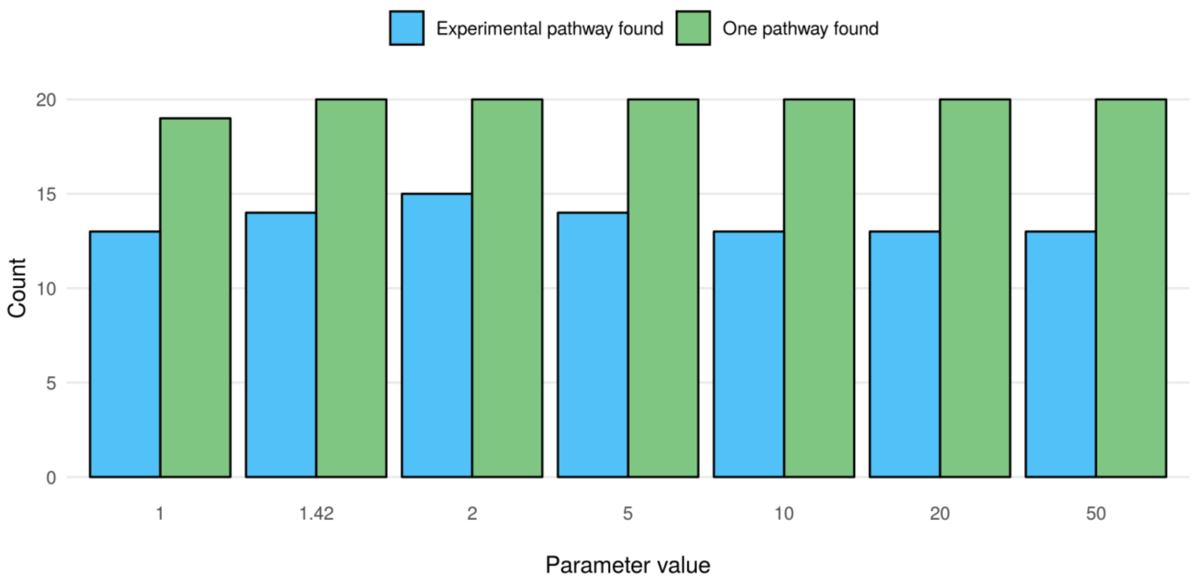


**Supplementary Figure 9: Impact of exploration constant value on retrieval performance of RetroPath RL.** We compared results between using different exploration constant values (UCTK). One pathway found means that at least one pathway has been predicted. Experimental pathway found means that the experimental pathway is from amongst the predicted pathways.

Virtual visits

This is the number of visits a new node starts with. The concept of virtual visits is that giving an initial value to a node will return more stable rollout results as they will be smoothed by a number less close to 0 rather than being very stochastic at low values. We can see this strategy did not give better results in our MCTS for bio-retrosynthesis in **Supplementary Figure 10**.

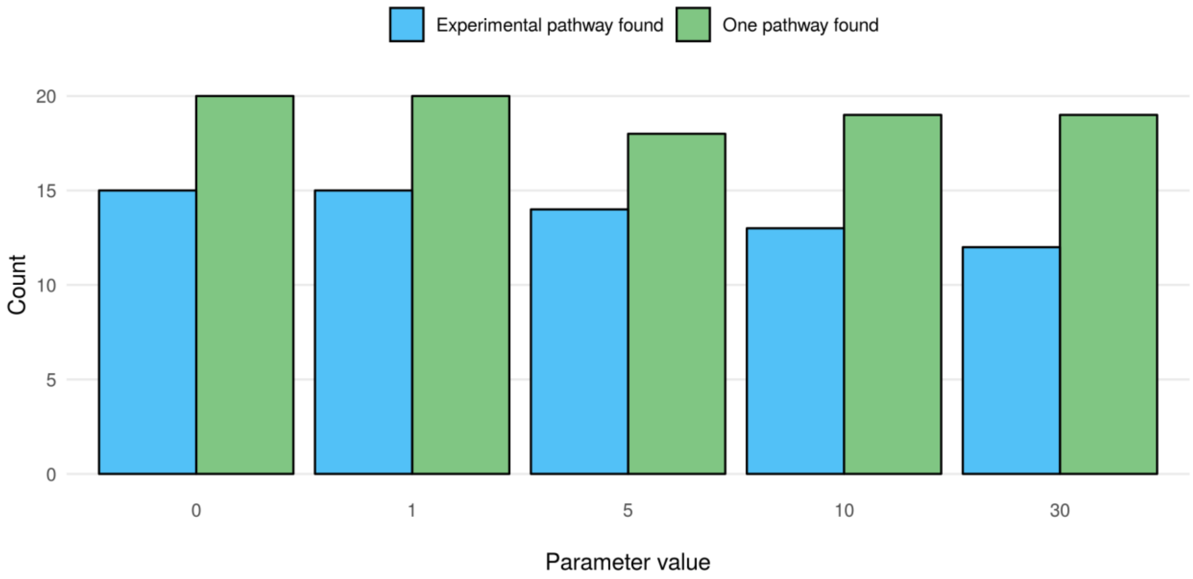


**Supplementary Figure 10: Impact of virtual visits on retrieval performance of RetroPath RL.** We compared results between using different virtual visits numbers. One pathway found means that at least one pathway has been predicted. Experimental pathway found means that the experimental pathway is from amongst the predicted pathways.

#### Solution rewarding

##### Penalty

This is the value returned when no compound of the state is within the chassis (including at the end of rollout). We can see in **Supplementary Figure 11** that increasing penalty does not yield better results in our case, and a value of -1 penalises enough the unsuccessful rollout results.

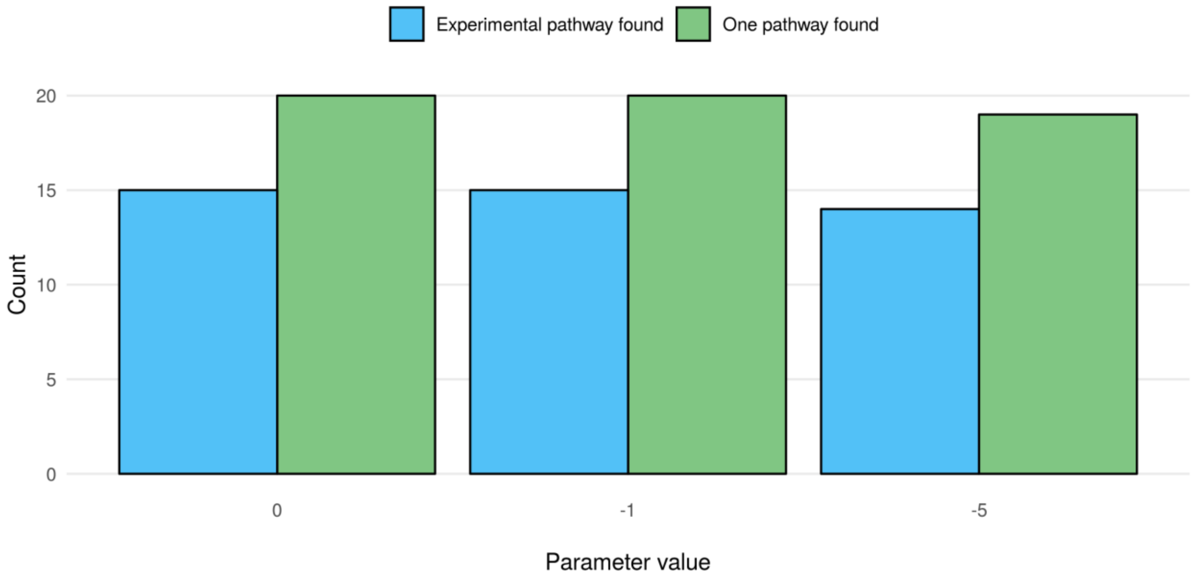


**Supplementary Figure 11: Impact of penalty on retrieval performance of RetroPath RL.** We compared results between using different penalties. One pathway found means that at least one pathway has been predicted. Experimental pathway found means that the experimental pathway is from amongst the predicted pathways.

##### Reward

Reward is the value returned when all compounds of the state are solved, to encourage exploration of the same area of the Tree. We can see from **Supplementary Figure 12** that a value of 5 provides a good trade-off between exploration of other areas of the tree and exploitation of promising regions.

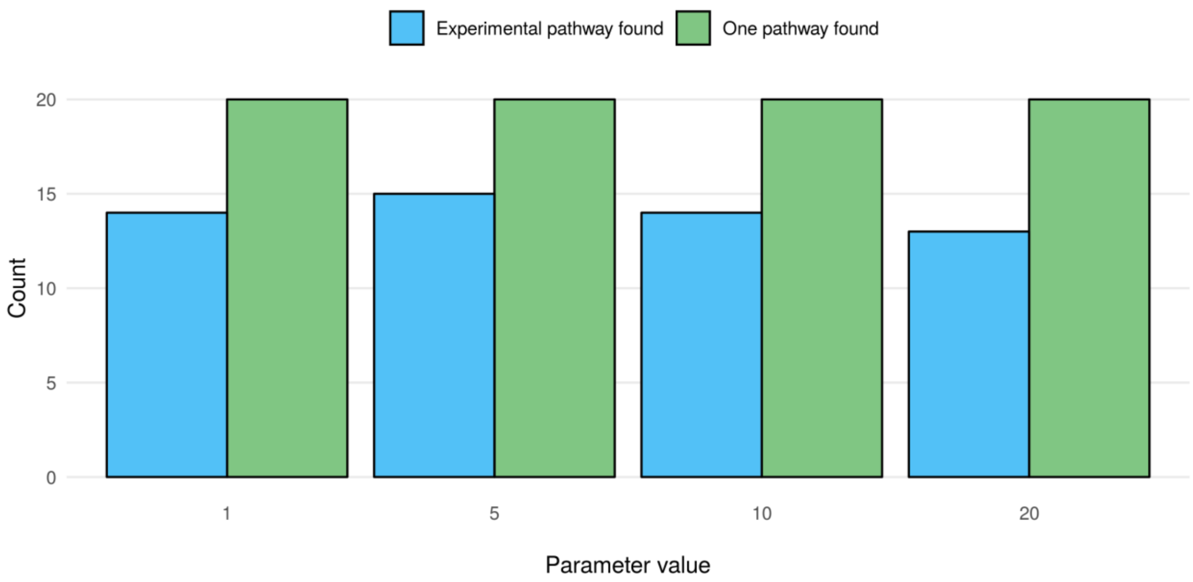


**Supplementary Figure 12: Impact of reward on retrieval performance of RetroPath RL.** We compared results between using different rewards. One pathway found means that at least one pathway has been predicted. Experimental pathway found means that the experimental pathway is from amongst the predicted pathways.

#### Other parameters - for other applications

Heavy saving

Saves search Tree state during the search instead of only at the end. Used to analyse Search evolution.

Stop at first result

The search stops once a single result is found.

Fire time-out and standardisation time-out

Timeouts on rule application on a compound.

Organism name

Choose another organism from our predefined list of sinks.

Complementary sink

Adding compounds to the sink as supplements. Can also be used to provide an entirely new sink following the required format.

#### Other parameters - exploratory

Remark: no detailed comparison was performed in this article on those parameters, contrary to the parameters presented above.

k_RAVE

For balancing of Rapid Action Value Estimation. The idea is to provide moves with results from rollouts elsewhere in the Tree, to give them an initial value. This will decrease in importance as the node itself is visited but provides a fast initial value.

Bias_k

When using bias (for example towards toxicity), how to weight this value.

Progressive bias

When used in conjunction with bias_k, can give an initial value to a node based on various policies: high reward, current state reward, no reward… This also helps initial estimation of the node value rather than rely only on costly rollouts.

Progressive widening

Allow a number of children different according to the number of visits of the node. This is to avoid expanding too much in spaces of the tree search that are actually not interesting.

#### Supplementary Note 2 : Detailed golden dataset analysis

Results comparing RetroPath RL and RetroPath 2.0 (run with the same set of rules at diameters 6, 10 and 16 and a timeout of 3 hours) on the golden dataset are presented in **Figure 2**. For all compounds, at least one pathway was found with those settings with RetroPath RL, while one compound had no pathway with RetroPath 2.0. For 2-amino-1,3-propanediol, the same core pathway was found, but the identified co-substrate in the first step was different (D-alanine predicted instead of D-glutamate for the experimental pathway, the main substrate being dihydroxyacetone phosphate). (one step different in **Figure 2** - dark blue). For four compounds (TPA, N-methylpyrrolinium, 1,4 BDO and protopanaxadiol), the experimental pathway was not found for different reasons. For TPA, the described experimental pathway in our golden dataset starts from a compound added to the mix, xylene. Running our workflow adding this compound to the sink allows us to find the experimental pathway (**Figure 2** - purple). For protopanaxadiol and N-methylpyrrolinium, we ran the MCTS using a different set of parameters, allowing both to explore more reactions (15 instead of 10) and more tolerance on the scores (cut-offs of 0.15 instead of 0.3). With those new settings we found the experimental pathway for both compounds. For 1,4 BDO, the experimental pathway was not found with these new settings either, but a similar pathway (lacking only one enzymatic step) was found with the default configuration. It transforms 4-hydroxybutyryl Coa into 1,4 BDO without using 4-Hydroxybutyraldehyde, supposedly catalysed by EC number (1.2.1.84: alcohol-forming fatty acyl-CoA reductase, without the alcohol dehydrogenase step from the literature example). The rest of this pathway is identical to the experimental pathway (one step lacking in **Figure 2** - light blue).

#### Supplementary Note 3 : Database speed-up calculations

When running a rule-based retrosynthetic algorithm, the most time and power consuming steps are rule application steps which require subgraph matching, as this is an NP hard subgraph isomorphism problem. We implemented a NoSQL database that allows for a frequent user to store rule application calculations. To allow for a fair comparison between runs and algorithms, it was not activated in the results presented previously. However, when this feature is active, the results of the first rule application on a compound is stored in the database. When the same calculation is encountered in a later run of RetroPath RL, results are retrieved from the database. This allows for faster runs of the algorithm and therefore larger and deeper exploration within the same time budget. For example, we ran the TPA retrosynthetic search 4 times: without the database, and with the database for the first, second and third time, allowing 1 hour and 100 000 iterations at each step. We present in **Supplementary Figure 13** the number of iterations performed in 1 hour, as well as the number of pathways found. While we can see the first run with the database is not as efficient as the run without it (reaches less iterations and does not find a pathway), we can see that having filled the database allows for more exploration of the tree in runs 2 and 3, where in the same allocated time more iterations are performed and more pathways found.

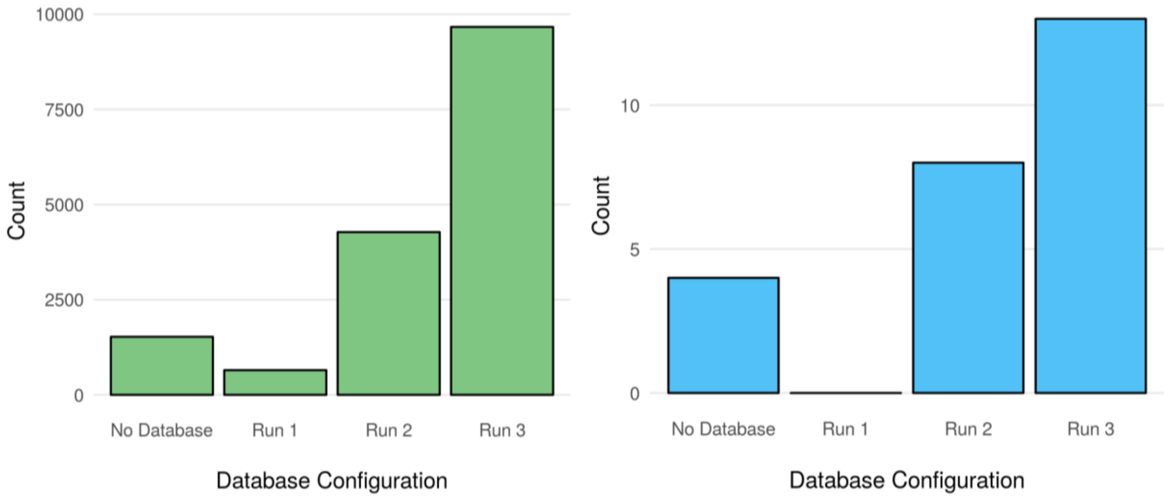


**Supplementary Figure 13: Database sped-up retrosynthetic search**. We compared results between a classical run where computation is performed on the fly versus storing results in a Database. **A)** Reached iteration is the number of iterations performed by the algorithm in the given timeframe (1hour). **B)** Found pathways is the number of found pathways per run.

*Method: Rule calculation cache using a NoSQL database*

Rule calculation can be optionally cached into a NoSQL database in order to optimise the running time of RetroPath RL. We released this cache system as an optional python package named “rp3_cache” that is available on GitHub at https://github.com/brsynth/rp3_cache. Technically, the cache system relies on the Mongo DB database (“Mongo DB” n.d.) that is embedded into a Docker (“Docker” n.d.) container to make the implementation agnostic of the operating system.

#### Supplementary Note 4 : Extending a previous search

Another feature of interest for expert users is the possibility to extend a previously run tree. For example, considering the example of protopanaxadiol, the experimental pathway was not found with our default settings, but was with more tolerant settings (15 children instead of 10 and a score cut-off of 0.15 instead of 0.3). However, instead of starting from scratch and losing the previously made calculations, it is possible to restart the search from a saved tree, with more tolerant settings. The results after running for 4h and 10 000 iterations are presented in **Supplementary Figure 14**. With the default settings, the iteration budget is spent finding only 1 pathway (not the experimental one). Using more tolerant settings is slower (604 iterations are performed in the allotted time) and 2 pathways are found. Extending the original tree allows for performing more iterations (786) and finding one more pathway, when compared to starting from scratch.

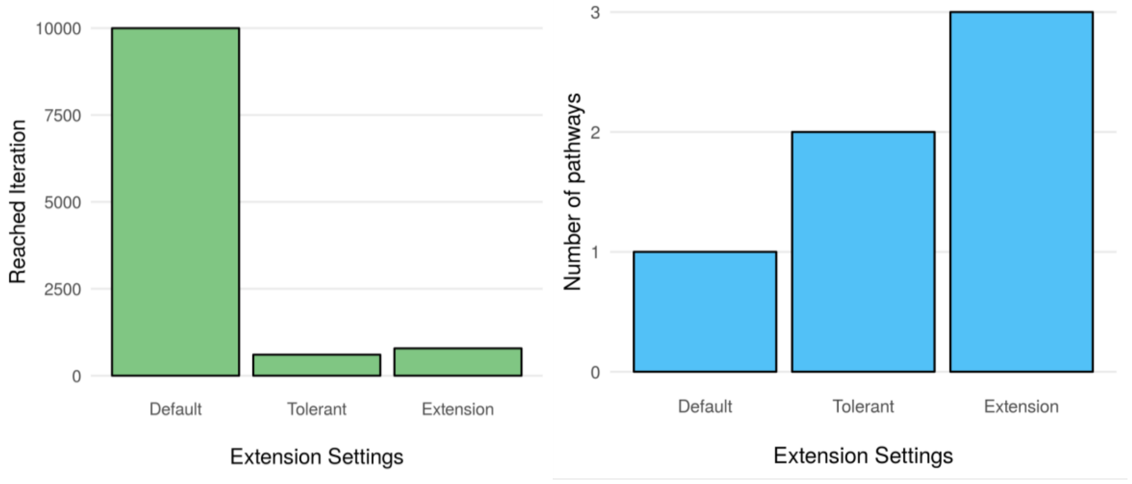


**Supplementary Figure 14: Extending a previous search**. We compared results between default settings, more tolerant settings and extending the saved searched. **A)** Reached iteration is the number of iterations performed by the algorithm in the given timeframe (1hour). **B)** Found pathways is the number of found pathways per run.

*Method: Tree extension*

The tree extension procedure starts with a tree containing search results. If n children were allowed in the first run and the extension allows m more children, up to n + m children can be found for a given node. Node scores and visit counts are re-initialised, and nodes are first flagged for extension then extended when they are first visited in the new search.

Yang, Jianming, and Lizhong Guo. 2014. "Biosynthesis of β-Carotene in Engineered E. Coli Using the MEP and MVA Pathways." *Microbial Cell Factories* 13 (November): 160.

Yim, Harry, Robert Haselbeck, Wei Niu, Catherine Pujol-Baxley, Anthony Burgard, Jeff Boldt, Julia Khandurina, et al. 2011. "Metabolic Engineering of Escherichia Coli for Direct Production of 1,4-Butanediol." *Nature Chemical Biology* 7 (7): 445–52.
